# A hidden integral structure endows Absolute Concentration Robust systems with resilience to dynamical concentration disturbances

**DOI:** 10.1101/830430

**Authors:** Daniele Cappelletti, Ankit Gupta, Mustafa Khammash

## Abstract

Biochemical systems that express certain chemical species of interest at the same level at any positive equilibrium are called “absolute concentration robust” (ACR). These species behave in a stable, predictable way, in the sense that their expression is robust with respect to sudden changes in the species concentration, regardless the new positive equilibrium reached by the system. Such a property has been proven to be fundamentally important in certain gene regulatory networks and signaling systems. In the present paper, we mathematically prove that a well-known class of ACR systems studied by Shinar and Feinberg in 2010 hides an internal integral structure. This structure confers these systems with a higher degree of robustness that what was previously unknown. In particular, disturbances much more general than sudden changes in the species concentrations can be rejected, and robust perfect adaptation is achieved. Significantly, we show that these properties are maintained when the system is interconnected with other chemical reaction networks. This key feature enables design of insulator devices that are able to buffer the loading effect from downstream systems - a crucial requirement for modular circuit design in synthetic biology.

## 1 Introduction

The network of chemical interactions of a biochemical system of interest can be complex and involve unknown reaction propensities. One of the main goal of reaction network theory consists in deriving dynamical properties from simpler graphical characteristic of the model, and independently on the specific value of kinetic parameters [22, 45]. The results presented in this paper follow this approach.

A qualitative property of great interest is the capability of a certain chemical species to be expressed with the same concentration at any positive steady state, independently on the initial conditions and on how many steady states are present. Namely, assume that the dynamics of the biochemical system are expressed by the *d*–dimensional ordinary differential equation

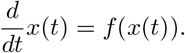

We say that the *i* – *th* species is *absolute concentration robust* (ACR), if there exists an *ACR value q* independent of the initial condition *x*(0) such that, whenever *x*(*t*) tends to a positive vector 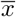, we have 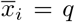. In the typical cases of interest, the positive steady state 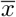 that is reached will depend on the initial condition *x*(0), while the entry 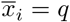 does not. As noted in [5], the property of absolute concentration robustness alone does not imply stability of the positive steady states: it only ensures that if a positive steady state 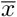 exists, then the value of the ACR species at 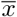 is the ACR value.

Under the assumption of stability, absolute concentration robustness provides a reliable, predictive response to environmental changes, since the species of interest reaches the equilibrium level relative to the new environmental setting, regardless the previous conditions. The existence and importance of this robustness property for various gene regulatory networks and signal transduction cascades is explored in many papers, including [1, 10–12, 35, 37, 39, 42, 43].

In the Control Theory community, and under the assumption of stability, the absolute concentration robustness is known as “robustness to disturbances in the initial conditions” [46, 47]. To achieve robustness with respect to some disturbance, the imbalance caused by the disturbance needs to be measured first. To this aim, a quantity of interest is the *integrator*, which is a function *ϕ* of the system variables whose derivative is exactly the imbalance to be eliminated. At steady state, the derivative of *ϕ* is zero and so needs to be the imbalance. In the setting of absolute concentration robustness, one would like to find an integrator whose derivative is the difference between the ACR species and its ACR value. Unfortunately, in general this cannot be done, as shown in Section 4.1 and as discussed in [46].

In the present paper, we systematically study for the first time the connection between ACR systems and integrators. Specifically, our first contribution is related to the existence of a linear combination of chemical species whose derivative is the difference between the ACR species and its ACR value, multiplied by a monomial. Such linear combination of species is called *constrained integrator* (CI), because it behaves similarly to an integrator given that the monomial does not vanish [46]. We rigorously prove that such a linear CI always exists for a large class of models that strictly includes the ACR systems introduced in [40]. This result has some important consequences: first of all, under the assumption of stability, it implies that the expression of ACR species is not only robust to changes in the initial conditions, but also to disturbances that are applied over time.

An important application in Synthetic Biology concerns the design of *insulators*. A number of biochemical systems are known to express a specific output if given a certain input. The systems can therefore be considered as modules with different functions. In cells, different modules are combined so that more complex responses to external stimuli become possible [30]. In Synthetic Biology, it is desirable to combine different modules to achieve the same level of complexity [38]. However, when connected, the different modules can affect the dynamics of each other and they can lose the desired dynamical properties they had when considered in isolation [17]. In a simplified framework, an *upstream* module processes an input, and its output is fed to a second, *downstream* module to be further processed. Since the information is passed in form of molecules, which are then consumed or temporarily sequestrated by the downstream module, the equations governing the upstream module dynamics are perturbed and its functionality can be affected. Such effect is commonly called *loading effect* [34] or *retroactivity* [17, 36], and needs to be minimized. In other words, the upstream module needs to be *insulated* from the loading effect caused by the downstream module. We propose two ways in which the robustness of the systems studied in this paper can be used to this aim. The first solution is to simply design an upstream module which is robust to loading effects, modeled as a persistent disturbance over time. The second solution is to design an extra component, called *insulator*, which transfers the signal from the upstream module to the downstream module while at the same time shielding the dynamics of the upstream module from retroactivity effects.

We will also show how more theoretical results on reaction network models can be obtained as a consequence of our work. In Reaction Network Theory, the study of steady state invariants consitute an interesting topic of research[18, 22, 33, 40]. In [40], it has been proven that certain graphical properties of the network imply the existence of an ACR species, regardless the choice of kinetic parameters. Such sufficient conditions are generalized in the present work while they are maintained simple to check. Moreover, no way to explicitly determine the ACR value was given in [40], and we fill the gap by proposing a fast linear method to calculate it. Furthermore, a substantial effort in the Reaction Network community is devoted to understand under what conditions dynamical properties of single systems can be lifted to larger systems [9, 24, 25, 29, 32]. Our contribution in this sense consist in proving that, under certain conditions, if an ACR system of the class studied in this paper is part of a larger model, the ACR species is still so in the lager system and its ACR value is maintained. Finally, it is worth mentioning that in the present work we consider the possibility of time-dependent rates for the occurrence of chemical transformations. This is more general than what is usually studied in Reaction Network Theory, with the exception of few works explicitly allowing for this scenario [2, 13, 14, 16, 28, 31].

## 2 Examples of ACR systems

### 2.1 An illustrative example

Consider two proteins *A* and *B*, whose interaction is described by

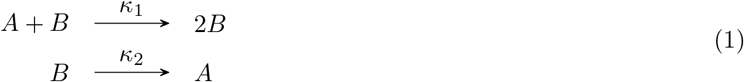

where the positive constants *κ*_1_ and *κ*_2_ describe the propensity of a reaction to occur. If enough proteins are present and they are homogeneously spread in space, then a good model for the time evolution of the concentrations of proteins *A* and *B* is given by mass-action kinetics. Specifically, the concentrations of *A* and *B* at time *t*, denoted by *x*_*A*_(*t*) and *x*_*B*_(*t*) respectively, are assumed to solve

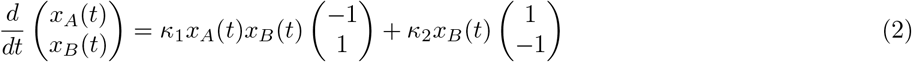

It is easy to check that the steady states of (2) are given by states (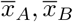) such that either 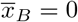 or 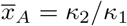. Hence, *A* is an ACR species because its expression at any positive steady state is the same. It is common during biochemical experiments to be able to control the inflow rate of some species (say *B*). Some additional chemical species may also be introduced, with the purpose of degrading some of the present components (in this case, species *C* is introduced to faster degrade species *B*). After these modifications, (1) becomes

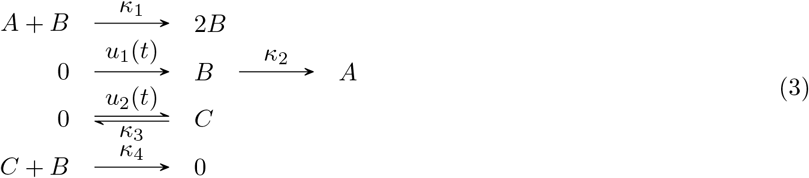

Since we still have

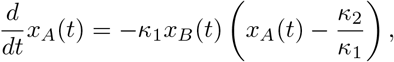

it is still true that the value of *x*_*A*_(*t*) will converge to *κ*_2_/*κ*_1_, as long as the functions *u*_1_, *u*_2_ and the rates *κ*_3_*, κ*_4_ are such that the species *B* is not removed from the system too fast. In this paper we will prove a general result describing when such kind of robustness to persistent perturbations is present for ACR systems.

### 2.2 EnvZ-OmpR osmoregulatory system

Consider the system described in Figure 1. The model is proposed in [40, 41] as osmoregulatory system in Escherichia Coli. It is in accordance with experimental observations discussed in [11, 37, 43]. According to the model, whose schematics is described in Figure 1, the activation rate of the sensor-transmitter protein EnvZ depends on the medium osmolarity. Then, an active form of EnvZ transfers its phosphoryl group to the sensory response protein OmpR, which becomes OmpR-P and promotes the production of the outer membrane porins OmpF and OmpC. Hence, it is important that the concentration of OmpR-P responds in a reliable, predictive way to changes in the medium osmolarity (which the rate constants *κ*_*i*_ in the reaction network of Figure 1 depend upon), but not on the initial concentration of the different chemical species involved. As a matter of fact, it follows from the results developed in [40] that OmpR-P is an ACR species.

**Figure 1:**
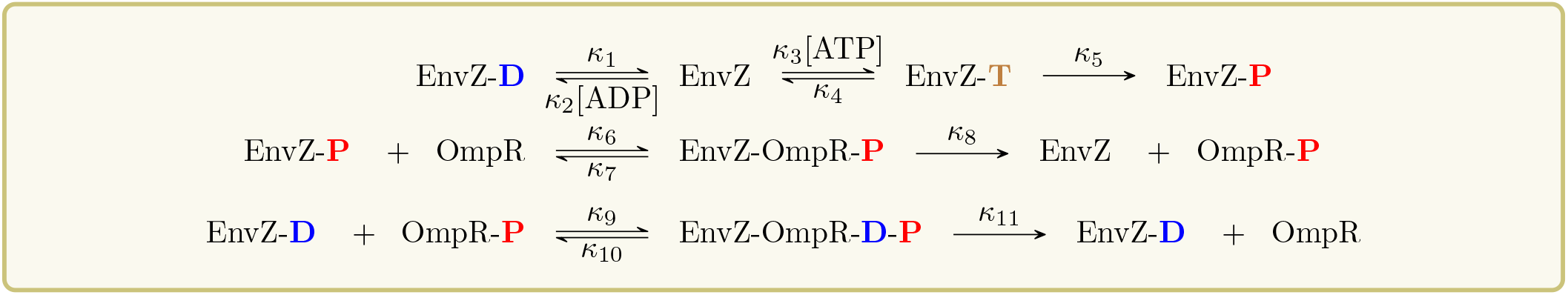
Proposed model for the EnvZ-OmpR signal transduction system in Escherichia Coli, which is able to explain the experimentally observed robustness in the expression of phosphorylated OmpR. In the first line of reactions, EnvZ can bind to ADP and ATP, but only when bound to ATP it can gain a phosphoryl group, and the resulting species is denoted by EnvZ-P. In the second line of reactions, EnvZ-P transfers the phosphoryl group to OmpR, through the formation of an intermediate complex. In the last line of reactions, the phosphoryl group is removed from OmpR-P through the action of EnvZ-D. The concentration of ATP and ADP is assumed to be maintained constant in time.

## 3 Necessary terminology and known results

In order to present the theory we develop, we first need to introduce some terminology. The linear combinations of chemical species appearing on either side of the chemical reactions of interest are called *complexes*, in accordance with the Reaction Network Theory literature. Be aware that the word “complex” has usually a different meaning in the Biology literature. We denote by *m* the number of complexes present in the network, an by *d* the number of chemical species. As an example, the complex of (1) are *A* + *B*, 2*B*, *B*, and *A*. Here, *d* = 2 and *m* = 4. In (3) the complexes are *A* + *B*, 2*B*, 0, *B*, *A*, *C*, and *C* + *B*, hence *d* = 3 and *m* = 7. Finally, in the system depicted in Figure 1 *d* = 8 and *m* = 10. Since a complex is a linear combination of species, each complex can be regarded as a vector of length *d*. For example, for the model (1) we can consider *A* + *B* as (1, 1), 2*B* as (0, 2), *B* as (0, 1), and finally *A* as (1, 0). With this in mind, we can define the *stoichiometric subspace* as

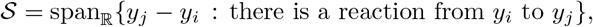

where *y*_*n*_ denotes the *n*th complex, for all 1 ≤ *n* ≤ *m*. For example, for (1) we have

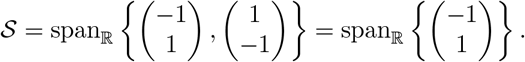

For (3), we have 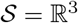.

In the most general formulation of reaction systems, a (time-dependent) rate function *λ*_*ij*_ is associated with the reaction from the *i*th to the *j*th complex of the network, and the concentration vector of the different chemical species is assumed to solve the differential equation

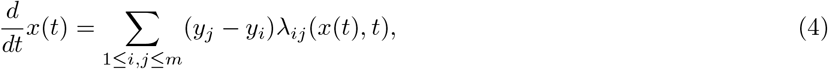

where if a reaction from the *i*th to the *j*th complex does not exist, then *λ*_*ij*_ is the zero function. Note that (4) simply sums the contributions to the dynamics given by the different chemical reactions. It is also not difficult to show that every solution to (4) is necessarily confined within a translation of the stoichiometric subspace. If for all non-zero propensities *λ*_*ij*_ there exists a positive constant *κ*_*ij*_ such that

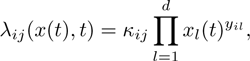

then the model is a *mass-action system*. In this case, (4) can be written as

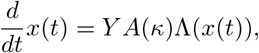

where *Y* is a *d* × *m* matrix whose *i*th column is *y*_*i*_, *A*(*κ*) is a *m* × *m* matrix given by

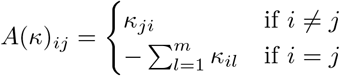

and Λ(*x*(*t*)) is a vector of length *m* whose *i*th entry is 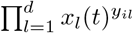. Examples of mass-action systems are (1) and the model in Figure 1.

A directed graph can be associated with a reaction network, where the nodes are given by the complexes and the directed edges are given by the reactions. Such a graph is called *reaction graph*. As an example, (1) is a reaction graph, while (3) is not because the complex 0 is repeated. The reaction graph corresponding to (3) is

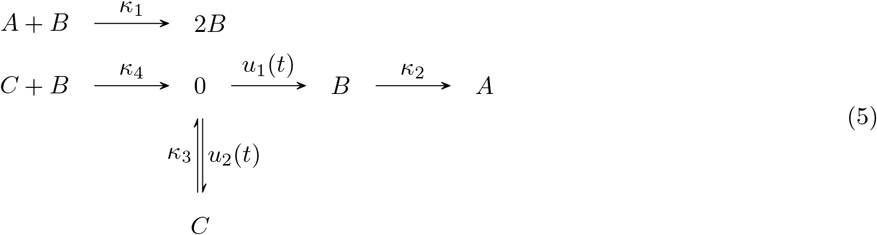

We denote by 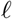 the number of connected components of the reaction graph associated with the network. For both the networks (1) and (3) 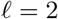, as well as for the EnvZ-OmpR osmoregulatory system of Figure 1. Then, we define the *deficiency* of a network as

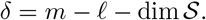

The deficiency of a network has important geometric interpretation, and a collection of classical deficiency theory results can be found in [21]. The deficiency of (1) is *δ* = 4 − 2 − 1 = 1, and the deficiency of (3) is *δ* = 7 − 2 − 3 = 2. Similarly, it can be checked that the deficiency of the EnvZ-OmpR osmoregulatory system in Figure 1 is 1.

Finally, we say that a complex *y* is *terminal* if for all paths in the reaction graph leading from *y* to another complex 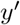, there is a path leading from 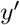 to *y*. If a complex is not terminal, then it is called *non-terminal*. As an example, the only terminal complexes for (1) and (3) are 2*B* and *A*.

We recall that a species is said to be *absolute concentration robust* (ACR) if its concentration at any positive steady state of (4) is the same. We are ready to state the following result, as presented in [40].

### Theorem 3.1.

*Consider a mass-action system, and assume the following holds:*

1. *there are two non-terminal complexes y_i_ and y_j_ such that only one entry of y_j_ − *y*_*i*_ is non-zero*;
2. *the deficiency is 1*.
3. *a positive steady state exists*.

*Then, the species relative to the non-zero entry of y*_*j*_ − *y*_*i*_ *is ACR*.

Note that a stronger version of Theorem 3.1 is proven in [40], which detects steady state invariant that are more general than the equilibrium concentration of a single species. The stronger version is stated in the Supplementary Material as Theorem B.3, and an extension of it is proven in the present work.

The model (1) has deficiency 1, as already observed, has at least one positive equilibrium and the non-terminal complexes *A* + *B* and *B* differ only for the species *A*. Hence, Theorem 3.1 applies and *A* is ACR. It is shown in [40] that the EnvZ-OmpR osmoregulatory system in Figure 1 also fulfils the hypothesis of Theorem 3.1, with the non-terminal complexes EnvZ-D and EnvZ-D+OmpR-P only differing for the species OmpR-P. As a consequence, OmpR-P is ACR. In Section 5 we will develop a method to explicitly calculate the ACR value through symbolic linear algebra. We note that Theorem 3.1 cannot be applied to (3) for two reasons: the model is not a mass-action system and its deficiency is 2.

As noted in [5], the positive steady states of a system with an ACR species are not necessarily stable. However, as a consequence of the present work (more precisely, as a consequence of Theorem 6.1 with *u* being the zero function), we know the following: if a mass-action system as in Theorem 3.1 has an unstable positive steady state, then either the system oscillates around it, or some chemical species is completely consumed, or some chemical species is indefinitely produced. We give here the formal definition of “oscillation”, as intended in this paper.

### Definition 3.1.

We say that a function *g* : ℝ_≥0_ → ℝ *oscillates* around a value 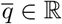 if for each *t* ∈ ℝ_≥0_ there exist *t*_+_ > *t* and *t*_−_ > *t* such that

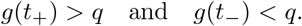

## 4 A linear constrained integrator

### 4.1 Control Theory background

In Control Theory, the focus is usually on systems of differential equations of the form

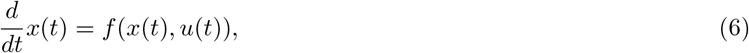

where 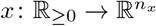 and 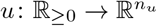 for some *n*_*x*_, *n*_*u*_ ∈ ℤ_>0_, and *f* is a differentiable function. The function *u* is called the *input of the system*. Further, a quantity of the form *z*(*t*) = *a*(*x*(*t*)) is of interest, where *a* is a differentiable function with 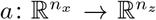, for some *n*_*z*_ ∈ ℤ_>0_. The function *z* is called the *output* of the system. In the usual setting, one needs to find an appropriate function *u* such that *z* is close to a desired level 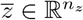, either on average or for *t* → ∞. To this aim, the existence of a function 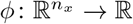 such that

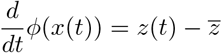

is of high importance, an is called an *integrator*. The name derives from *ϕ*(*x*(*t*)) being the integral of the error that needs to be controlled:

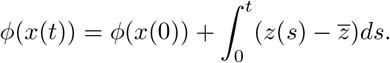

If the function is fed back to the system and is used to tune the input, then an *integral action* or *integral feedback* is in place [8, 19]. One of the main features of an integrator is that the derivative of *ϕ*(*x*(*t*)) is zero if and only if 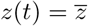. If a function 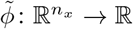 satisfies

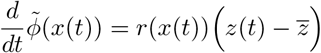

for some differentiable function 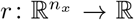, then 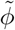 is called a *constrained integrator* (CI) [46]. The name derives from the fact that the derivative of 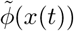 is zero if and only if 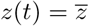, provided that *r*(*x*(*t*)) ≠ 0. In biology, it is common to find CIs, and the condition *r*(*x*(*t*)) ≠ 0 is usually implied by *x*(*t*) ≠ 0 [46]. Note that in [46] an explicit distinction between integrators and integral feedbacks is not made.

In the setting of systems with ACR species, the output *z* can be considered to be the concentration of the ACR species over time, and 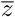 can be their ACR values. In (1), *z*(*t*) = *x_A_*(*t*) and 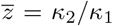. A CI (as noted in [46]) is given by 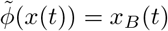, since

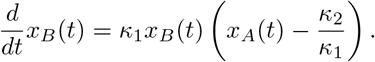

The question of whether an integrator exists can be quickly answered in negative, because any point of the form 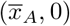 is a steady state. If an integrator *ϕ* existed, then by choosing 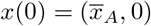 we would have

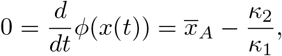

which cannot hold expect for a specific value of 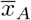. An integrator may still exist in a weaker sense, if we restrict its domain. For example, in this case the function 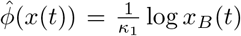 would be an integrator, in the sense that if *x*_*B*_(*t*) > 0 then

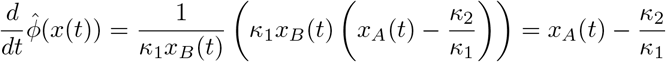

However, the domain of 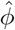 is not the entire ℝ^2^. Finally, since linear functions could always be extended continuously to the boundaries of 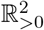, a linear integrator cannot exist for (1) even if its domain is restricted.

### 4.2 Existence and characterization

We state here our result concerning linear CIs. A stronger version is proved in the Supplementary Material. The result is inspired by the analysis carried on in [40], which is here expanded.

For any *n* × *l* real matrix *M* and real vector *v* of length *n*, we denote by (*M|v*) the *n* × *l* + 1 matrix obtained by adding the column *v* at the right of the matrix *A*. Let 1 ≤ *i*, *j* ≤ *m*. To present our result, we first need to define the preimage

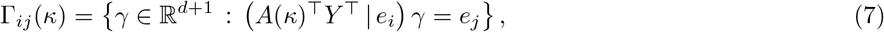

where *e*_*n*_ denotes the *n*th vector in the canonical basis of ℝ^*m*^, whose *n*th component is 1 and whose other components are 0. The role of Γ_*ij*_(*κ*) is that of providing vectors *γ* satisfying

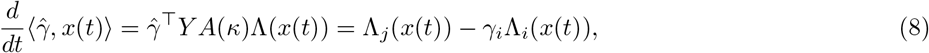

where 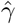 is the projection onto the first *d* components of *γ*, and 〈·, ·〉 is the standard scalar product. Under certain assumptions, (8) will provide us with a CI. Then, the projection of Γ_*ij*_(*κ*) onto the first *d* coordinates will be of interest, and we will denote it by 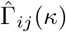. The set Γ_*ij*_(*κ*) can be calculated with symbolic linear algebra. We will also prove in the Supplementary Material how 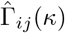 is connected with 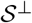, which is a set easily described by linear algebra and independent on the rate functions. Specifically, we will prove that if 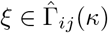, then necessarily

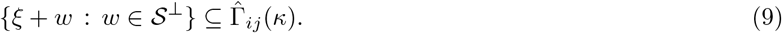

We will also give sufficient conditions under which the inclusion in (9) is an equality.

As an example, consider the model in Figure 1. Using Matlab, we quickly obtain that a vector *ξ* is in Γ_18_(*κ*), with

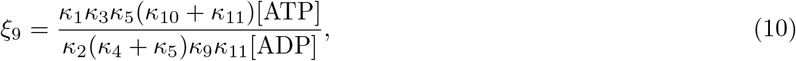

and it is shown in the Supplementary Material that

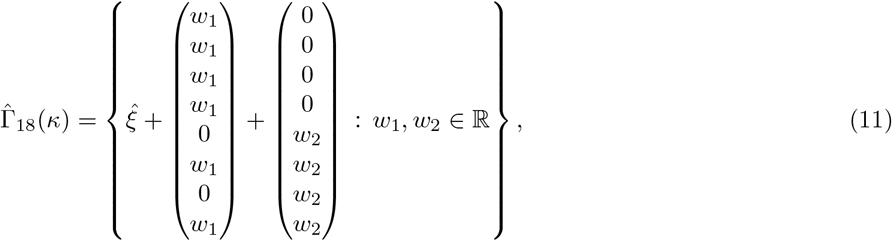

where 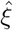 is the projection of *ξ* onto its first *d* = 8 coordinates.

The family of models we study in this paper concerns reaction systems with two non-terminal complexes *y*_*i*_ and *y*_*j*_ differing in just one entry, for which 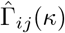 is non-empty. Our first result shows that such a family includes the models studied in [40]. The proof can be found in the Supplementary Material.

#### Theorem 4.1.

*Consider a mass-action system, and assume the following holds*:

1. *there are two non-terminal complexes y*_*i*_ *and y*_*j*_ *such that only one entry of y*_*j*_ − *y*_*i*_ *is non-zero*;
2. *the deficiency is 1*.
3. *a positive steady state exists*.

*Then*, 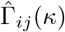 *is non-empty*.

As an example, we know already from direct calculation that for the EnvZ-OmpR signaling system (11) holds, which in turn implies that the set 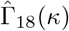 is non-empty. However, we could have also derived this information from Theorem 4.1, without explsicitly calculating 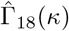.

We note here that the converse of Theorem 4.1 does not hold. We show this with an example of a multisite phosphorylation signaling system in the Supplementary Material, which does not fall in the setting of [40] but for which we are able to prove absolute concentration robustness regardless the choice of rate constants, as long as a positive steady state exists. Notably, we are also able to derive information on when this occurs without working directly with the differential equation. As a consequence of this example, the family of models we analyze is proven to be strictly larger than that studied in [40]. The following holds.

#### Theorem 4.2.

*Consider a mass-action system. Assume that there are two complexes y*_*i*_ *and y*_*j*_ *only differing in the nth entry, and that* 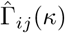 *is non-empty. Let* 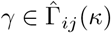, *and define*

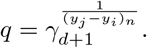

*Then, either no positive steady state exists or the nth species is ACR with ACR value q. Moreover*,

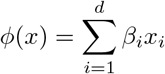

*is a linear CI with*

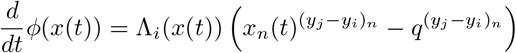

*for any initial condition x*(0) *if and only if* 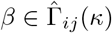.

The existence of a linear CI given by Theorem 4.2 is essential to develop the results presented in the next sections. Before unveiling the consequences of Theorem 4.2, however, it is important to stress that a CI does not necessarily constitute a feedback, as one may be tempted to think. Consider

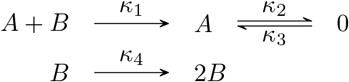

with *κ*_1_*κ*_3_ = *κ*_2_*κ*_4_. It can be shown that the system satisfies the conditions of Theorems 3.1 and 4.1, with the non-terminal complexes *A* + *B* and *A* differing only in species *A*. Hence, *A* is ACR and the assumptions of Theorem 4.2 hold. A linear CI as in Theorem 4.2 is given by *ϕ*(*x*) = −*x*_*B*_/*κ*_1_, since for this choice

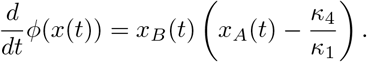

However, the quantity *ϕ*(*x*(*t*)) does not regulate the dynamics of *A*, since

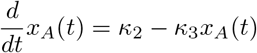

does not depend on *x*_*B*_(*t*). Since in this case the CI is not acting on the system, it is not surprising that the existence of positive steady states is lost as soon as *κ*_1_*κ*_3_ ≠ *κ*_2_*κ*_4_.

It is also worth mentioning that not all systems with ACR species have a linear CI: consider the mass-action system

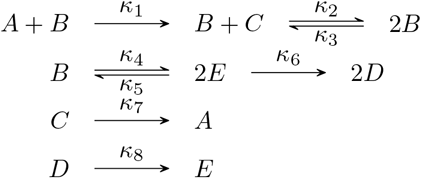

The model is considered in [3], where it is proven that the species *A* is ACR. We show in the Supplementary Material that there exists no linear function *ϕ* whose derivative at time *t* is of the form *r*(*x*(*t*))(*x*_*A*_(*t*)^*γ*^ − *q*), for some polynomial *r* and some real numbers *γ, q*. Note that in this case Theorem 4.1 does not apply because the deficiency of the network is 2.

## 5 A method to calculate the ACR value

The first interesting consequence of Theorem 4.2 is that the ACR values of the mass-action systems satisfying the assumption of the theorem can be calculated by finding at least one element of the preimage Γ_*ij*_(*κ*), and this can be done via a simple symbolic linear algebra calculation. As an example, consider the EnvZ-OmpR osmoregulatory system in Figure 1. Then, Theorem 4.2 implies that the ACR value of OmpR-P is the value given in (10). This value is in accordance with the one found in the Supplementary Material of [40], however we found it by calculating a single element in the preimage of a matrix, as opposed to working with the rather complicated differential equation associated with the model. An even more involved examples is dealt with in the Supplementary Material.

## 6 Rejection of persistent disturbances

### 6.1 The result

We state an important consequence of Theorem 4.2, a stronger version of which is proven in the Supplementary Material:

#### Theorem 6.1.

*Consider a mass-action system, with associated differential equation*

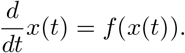

*Assume that there are two complexes y_i_ and y*_*j*_ *only differing in the nth entry, and that* 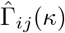 *is non-empty. Let q be the ACR value of the nth species. Consider an arbitrary function u with image in* ℝ^*d*^ *such that a solution to*

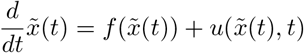

*exists. Assume that there exists a* 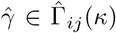 *which is orthogonal to the vector u*(*x, t*) *for any x, t. Then, for any initial condition* 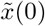, *at least one of the following holds:*

a. *the concentration of some species goes to 0 or infinity, along a sequence of times;*
b. 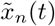 *oscillates around q and* 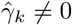 *for some k* ≠ *n*;
c. *the integral*

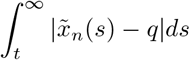

*tends to* 0, *as t goes to infinity*.

The result implies that if a disturbance orthogonal to a vector 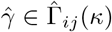 is applied over time, then the stability of the ACR species is maintained: at most, the ACR species can be forced to oscillate around its original ACR value, but it cannot be forced to attain another equilibrium level without causing extinction or overexpression of the chemical species present. We analyze the power of Theorem 6.1 by showing some examples of applications.

*Example* 6.1. Consider the mass-action system (1), which fulfills the assumptions of Theorem 6.1 as already observed. Assume the complexes are ordered as *A* + *B*, 2*B*, *B*, and *A*, and the species are ordered alphabetically as *A*, *B*. Hence, the two non-terminal complexes differing in the ACR species *A* are the 1st and the 3rd, and it is shown in the Supplementary Material that

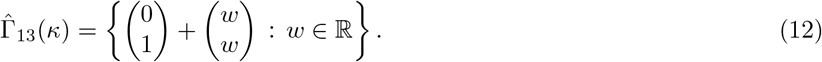

Hence, by choosing *w* = −1, we have that

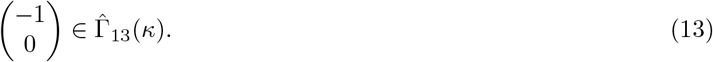

This vector is clearly orthogonal to any disturbance acting on the production and degradation rates of the species *B*. Hence, it follows that the stability and the ACR value of the species *A* is maintained in (3), provided that no species is completely removed or indefinitely expressed. Specifically, since the entry of (13) relative to *B* is zero, it follows from Theorem 6.1 that if all the species concentrations are bounded from below and from above by positive quantities, necessarily the concentration of the species *A* converges to its ACR value as *t* goes to infinity, despite the disturbances.

*Example* 6.2 (EnvZ-OmpR osmoregulatory system). Consider the osmoregulatory system in Figure 1, whose features have already been discussed in the paper. In particular, we know the species OmpR-P is ACR with ACR value (10). Recall that we ordered the complexes such that the two non-terminal ones differing in OmpR-P are the 1st and the 8th. It follows from (11) that for any chemical species, there is a vector in 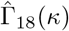 with the associated entry equal to 0. It follows that even if the production and degradation of any chemical species in the model is tampered with, the stability and the ACR value of the species OmpR-P are maintained, in the sense described by Theorem 6.1.

We can push the disturbances further. By appropriately choosing *w*_1_ and *w*_2_ in (11), we can see that there is vector in 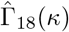 whose entries relative to the species OmpR and EnvZ are both 0. Hence, it follows by Theorem 6.1 that by tampering with the production and degradation rates of both these species over time, if no extinction and no overexpression occurs, then the concentration of OmpR-P still converges to the value (10), or oscillates around it.

As a final remark, we note that (9) can be useful in determining whether a vector in Γ_*ij*_(*κ*) exists, with a specific component equal to 0, say the *n*th one. In fact, the existence of such a vector can be deduced without calculations, if there is a vector 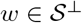 whose *n*th component is different from 0.

### 6.2 Insulating properties

Here we illustrate how the theory we developed can be utilized to solve the Synthetic Biology problem of retroactivity. As explained in the Introduction, the loading effects caused by a downstream biochemical module can disrupt the functionality of upstream modules, which prevents the implementation of biochemical circuits by interconnecting biochemical modules with different functions [17]. A concrete example of loading effect is illustrated in Figure 4.

Assume that a mass-action system has two complexes *y*_*i*_ and *y*_*j*_, that are only different in the *n*th component, which corresponds to the species *X*. Assume further that 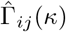 is non-empty and that a positive steady state exists. Hence, the species *X* is ACR, with some ACR value *q*. It further follows from Theorem 6.1 that, if there exists 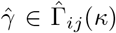 with 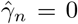, then the production and degradation rates of the species *X* can be arbitrarily perturbed over time by an arbitrary function *u*, without compromising its robustness. Specifically, if the perturbed system is stable and no chemical species is completely consumed, then the concentration of *X* will still converge to the same ACR value *q* as in the original mass-action system. The key observation we make here is that the perturbation *u* can be considered as the loading effect of a downstream module that takes the concentration of species *X* as input. In this case, the loading effect on the original mass-action is rejected and the concentration of *X* is maintained at a desired level *q* at steady state. Further, the concentration of *X* is maintained approximately constant in the transient dynamics as well, if we assume as done in [17] that a separation of dynamics time scale is in place. Specifically, assume

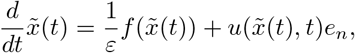

for some small *ε* > 0, with 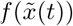 and 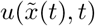 being of the same order of magnitude. Under the assumption of stability, if *ε* is very small then the perturbed system will quickly approach the slow manifold defined by

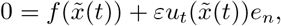

where *u*_*t*_ is a function from ℝ^*d*^ to ℝ^*d*^ defined by *u*_*t*_(*x*) = *u*(*x*, *t*). By Theorem 6.1 applied to the disturbance *u*_*t*_, the species *X* assumes its ACR value at any positive point of the slow manifold, which is exactly what we wanted.

As an example of application, consider the EnvZ-OmpR osmoregulatory system in Figure 1. It follows from (11) that there exists 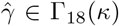 with a zero in the entry corresponding to the ACR species OmpR-P. Hence, the production and degradation rates of OmpR-P can be arbitrarily changed over time, without altering its robustness property, in the sense described by Theorem 6.1. As observed, the statement still holds true if the perturbation is originated by a downstream module that acts on OmpR-P. Hence, the EnvZ-OmpR osmoregulatory system can be used to maintain the expression of OmpR-P at a desired level, which depends on the input rate constants, even if the species OmpR-P is used by a downstream module. Moreover, if the downstream module acts on a slower time scale, the concentration of the species OmpR-P is approximately maintained at the target level at any time point. In Figure 2, a diagram describing this situation is proposed.

Consider now the case were an upstream module is affected from loading effects. We show how the theory developed in this paper can be used to design an insulator. Assume that the upstream module accepts *u*(*t*) as input, and modulates the concentration of the species *A*\* accordingly. The species *A*\* is then used by a downstream module, which returns a function of the concentration of *A*\* as output. The action of the downstream module on the species *A*\* causes a loading effect on the upstream module. This loading effect should be reduced. To this aim, we proposed to modify the downstream module such that it acts on a species *A* rather than on the species *A*\*, and to include in the system the following module, where *B* is a species that is not used by neither the upstream nor the downstream module:

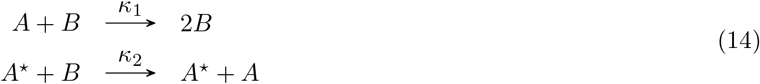

**Figure 2:**
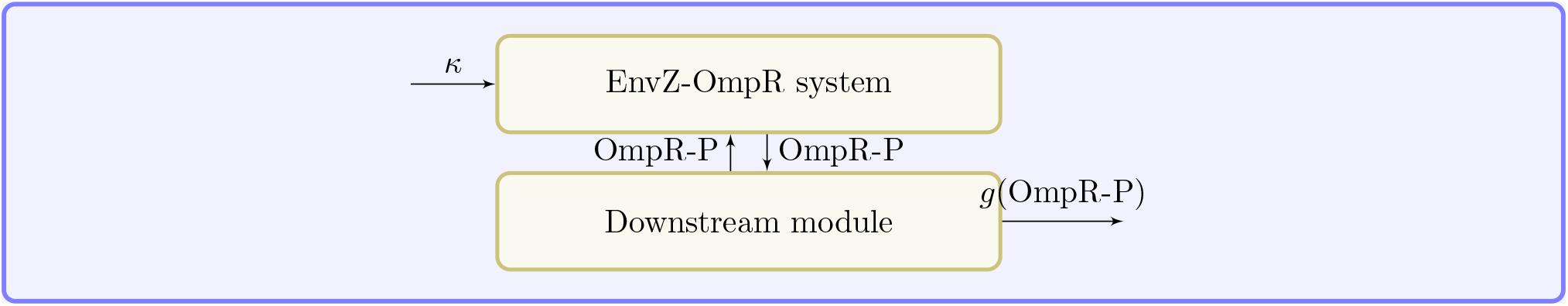
Proposed use of the EnvZ-OmpR signal transduction system of Figure 1 as a controller of a downstream module utilizing OmpR-P. The concentration of OmpR-P is regulated by the EnvZ-OmpR signaling system, with equilibrium given by (10). The equilibrium can be adjusted by modifying the parameters *κ*_*ij*_ of the EnvZ-OmpR signaling system (which depend on the medium osmolarity) or the ratio between ADP and ATP present. The output of the downstream module is a function *g* of the concentration of OmpR-P, which is received as input.

Assume stability is reached and that the species *B* is not completely consumed. Then, at steady state the concen-tration level of *A\** is fixed, and the concentration of the ACR species *A* will converge to its ACR value *κ*_2_*x*_*A*_* /*κ*_1_ regardless any disturbance applied to the production and degradation rate of *A*. In fact, a linear CI as in the statement of Theorem 4.2 is given by *ϕ*(*x*) = *x*_*B*_/*κ*_1_, and at any time point

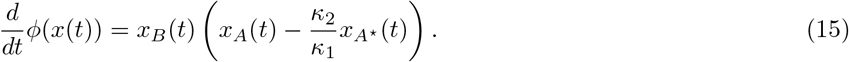

We further note that if the dynamics of (14) occur on a faster time scale than the rest of the system, then a slow manifold is quickly approached were the concentration of the species *A* is maintained at the level *κ*_2_*x*_*A\**_ (*t*)/*κ*_1_ at any time point. In this case, the module (14) approximately outputs a multiple of the concentration of *A*\* over the whole time line. The multiplicative constant can be tuned through the parameters *κ*_1_ and *κ*_2_, as well as the time scale that (14) operates in. The time scale can be further tuned via the concentration of *B*, as it also follows from (15). In conclusion, the downstream module receives as input a good approximation of a multiple of the concentration of *A*\*, and its activity does not affect the upstream module, nor (14). Moreover, (14) does not affect the upstream module at all, since the species *A* appears in (14) as a catalyst and is not changed in the catalysed reaction. The proposed insulating strategy is illustrated in Figure 3, and it is applied to an example discussed in [17] in Figure 4.

**Figure 3:**
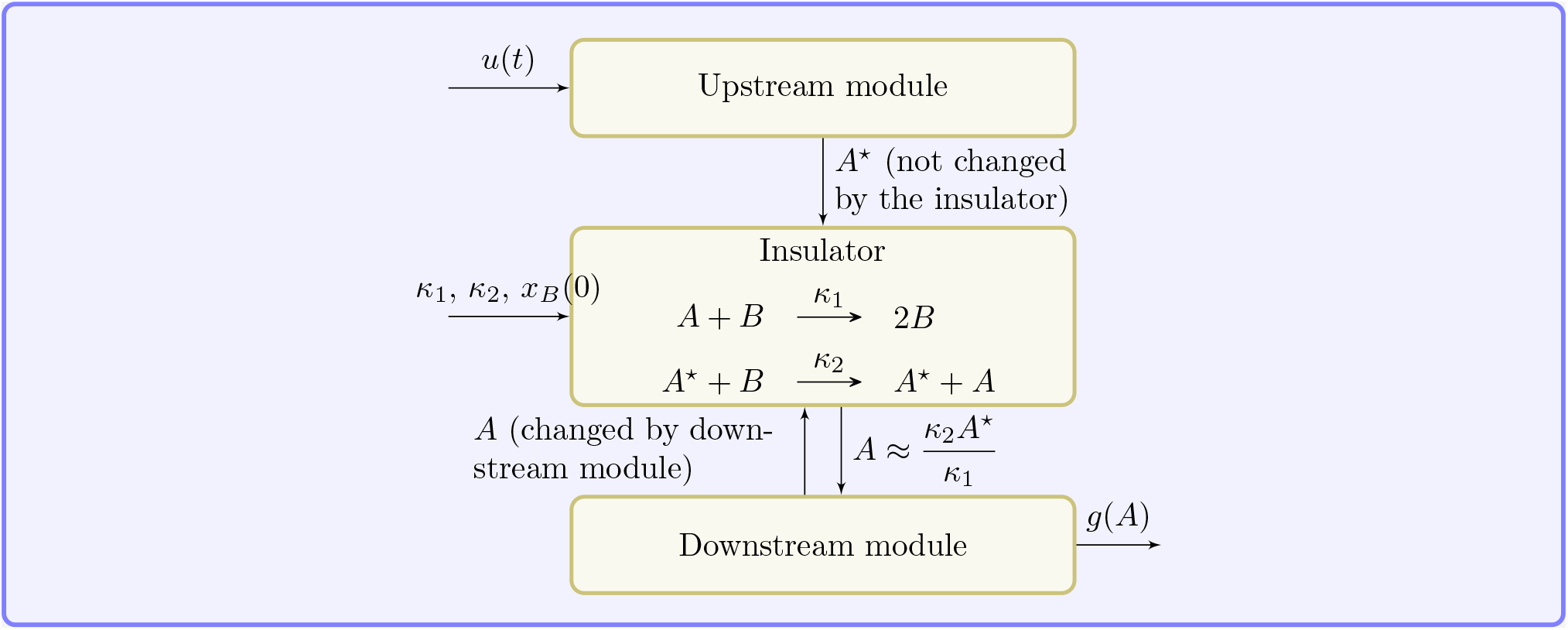
The upstream module expresses the chemical species *A*\* as output. The insulator transfers a multiple of the signal from the upstream module to the downstream module, which is modified to accept as input the concentration of *A* rather than the concentration of *A*\*.

**Figure 4:**
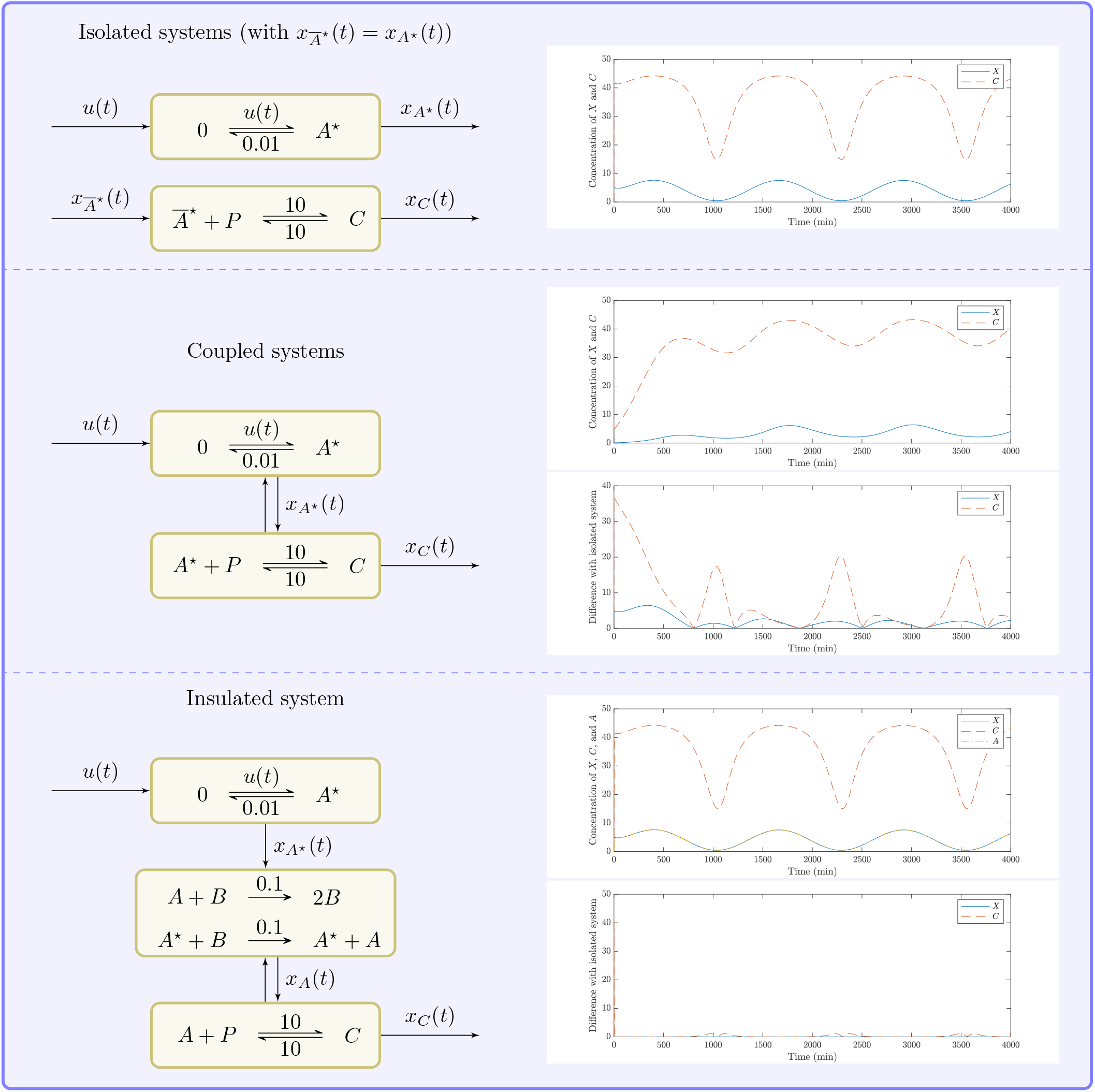
In the example, we let *u*(*t*) = 0.04(1 + sin(0.005*t*)) and *x_A*_* (0) = 5. All the rates are in 1*/min*. The ODE solution is calculated in Matlab with `ode23s`. In the first panel, the two systems are considered in isolation, with the assumption that the concentration of the species 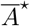 is maintained at the same level as the concentration of species *A*\*, at any time point. In the second panel, the two systems are linked together, and the species *A*\* is directly used by the downstream system. The dynamics are completely disrupted by the loading effects, a plot of the absolute value of the difference of the two solutions over time is proposed. In the third panel, the insulator of Figure 3 is utilized, with *x*_*A*_(0) = 0 and *x*_*B*_(0) = 20. The loading effect are practically removed, despite the choice of low rate constants 0.1. Specifically, the difference of the concentration of *C* between the solution of insulated system and the solution of the isolated systems spikes quickly to 40, but it decreases to less than 0.5 within 10 minutes, after which is maintained low as illustrated in the second plot of the third panel.

### 6.3 Inclusion in larger systems

In the previous section, we have seen how the stability properties of an ACR system can be transferred, in a sense, to a larger model including further chemical transformations and external inputs. As a particular case, Here, we explicitly state when the property of absolute concentration robustness can be lifted from portions of the biochemical system to the whole system. As a consequence, we further extend the set of sufficient conditions of Theorem 3.1 that imply the existence of an ACR species. Before stating the relevant result, which is a consequence of Theorem 4.2, we need a definition. Given a reaction system 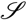, we say that 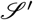 is a sub-system of 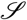 if it can be obtained from 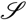 by canceling some reactions, and if the choice of rate functions for the remaining reactions is maintained. Moreover, if 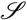 and 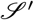 have *d* and 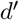 species, respectively, we let 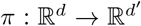 be the projection onto the species of 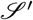. The following holds:

#### Corollary 6.2.

*Consider a reaction system 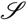, and let 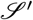 be a sub-system. Assume that 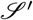 is a mass-action system with two complexes π(y_i_) and π(y_j_) only differing in the entry relative to the species X, and for which 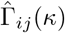 is non-empty. Moreover, assume there exists 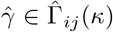 such that π(y_l_ y_k_) is orthogonal to 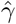 for all y_k_ ⟶ y_l_ that are reactions of 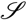 but not reactions of 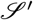. Then, the X is ACR for both 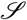 and 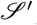, with the same ACR value.*

The proof of a stronger result is in the Supplementary Material. Here we illustrate how the corollary can be applied in the case of EnvZ-OmpR signaling system: consider the reaction system 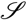 described in Figure 5, which includes the EnvZ-OmpR osmoregulation system described in Figure 1 as a sub-system. We assume that a protein can misfold when the phosphoryl group is transferred from EnvZ to OmpR. Such misfold can be corrected by chaperons, which are independently produced and degraded through a mechanism which we assume unkown, but that does not involve EnvZ or OmpR proteins. We also allow for an arbitrary and persistent external control on the expression level of EnvZ sensor-transmitter protein. Finally, we consider the utilization of OmpR-P as transcription regulatory protein of the outer membrane porins OmpF and OmpC. For our purposes, we assume the details of the transcription mechanism are not known, but that only the protein Ompr-P is involved in the process. As previously done, let the complexes of the EnvZ-OmpR osmoregulation system be ordered from left to right and from top to bottom, such that EnvZ-D and EnvZ-D+OmpR-P are the 1st and the 8th complex, respectively. Also, let the species be ordered according to their appearance from left to right and from top to bottom, as EnvZ-D, EnvZ, EnvZ-T, EnvZ-P, OmpR, EnvZ-OmpR-P, OmpR-P, EnvZ-OmpR-D-P. In particular, Envz is the second species, EnvZ-OmpR-P is the sixth species, and OmprR-P is the seventh species. It follows from (11) that, by choosing *w*_1_ = −*ξ*_2_ and *w*_2_ = −*ξ*_7_ = 0, a vector 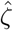 is in 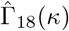 with:

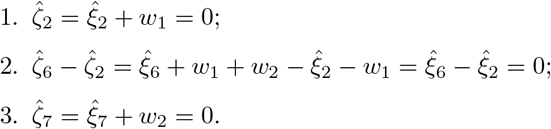

**Figure 5:**
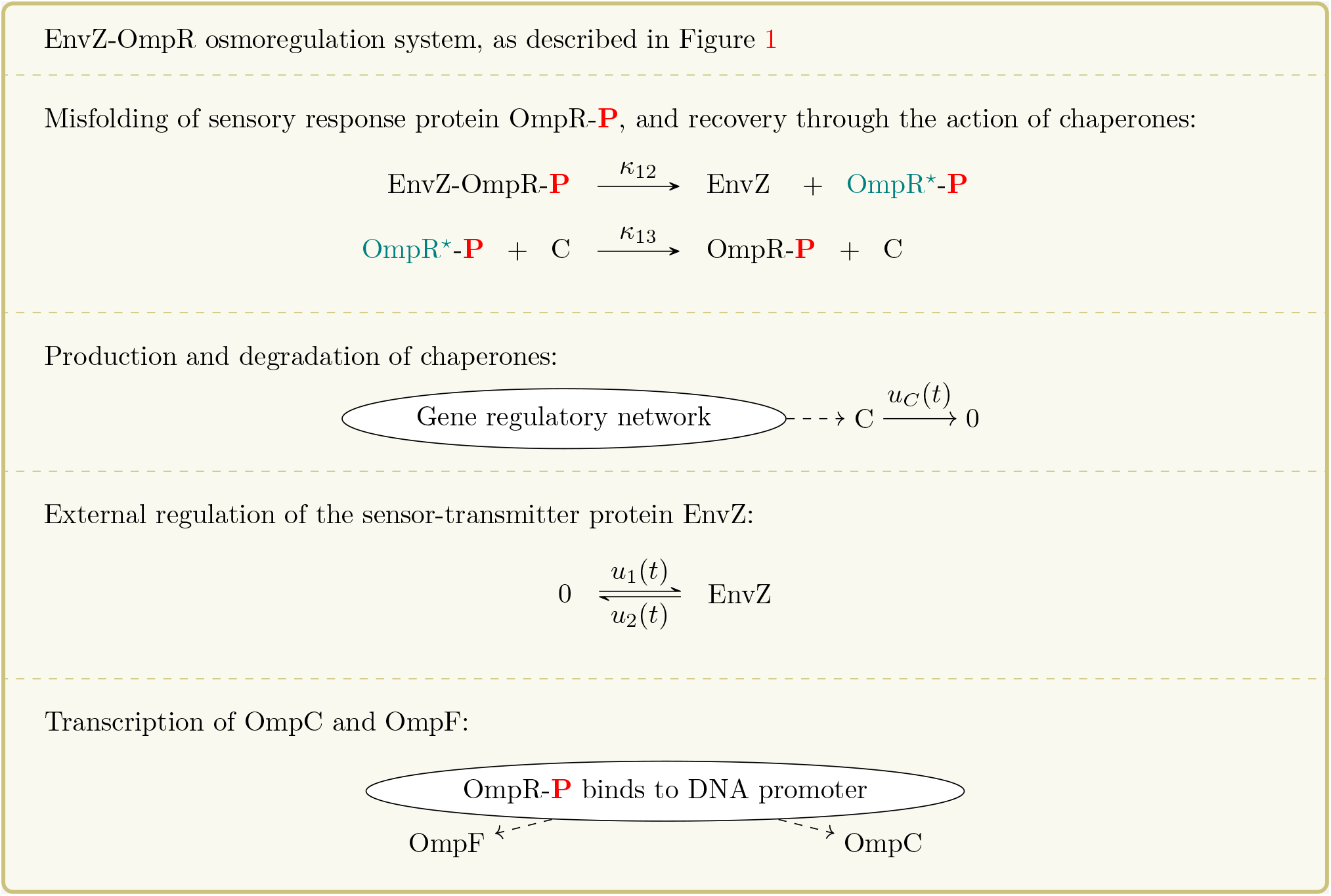
Reaction system including the EnvZ-OmpR osmoregulation system depicted in Figure 1 as a sub-system. Parts of the system depicted here are unknown, specifically no model is given for the production of chaperons or for the transcription of the outer membrane porins OmpF and OmpC.

Denote by *E*_*k*_ the vector of ℝ^*d*^ with the *k*th entry equal to 1 and the other entries equal to zero. The following holds.

*Misfolding of OmpR-P*. The projection of the difference between EnvZ + OmpR*-P and EnvZ-OmpR-P onto the species of the EnvZ-OmpR signaling system is *E*_2_ − *E*_6_, which is orthogonal to 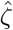 by 2. The projection of the difference between OmpR-P + C and OmpR*-P + C is *E*_7_, which is orthogonal to 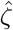 by 3.

*Production and degradation of chaperons*. By assumption, any chemical reaction 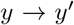 involved in the production and degradation of chaperones does not consume or produce any chemical species of the EnvZ-OmpR signaling system. Hence, 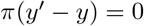, which is orthogonal to 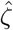.

*External regulation of EnvZ*. The difference between EnvZ and 0 is ±*E*_2_, which is orthogonal to 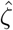 by 1

*Transcription of OmpC and OmpF*. We assume that the transcription only involves the species OmpR-P, out of all the species in the EnvZ-OmpR osmoregulation system. Hence, for all the reactions 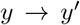 involved in the transcription, either 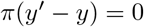 or 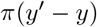 is a multiple of *E*_7_. In either case, 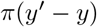 is orthogonal to 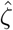.

It follows from Corollary 6.2 that the species OmpR-P is ACR in the reaction system of Figure 5. Moreover, its ACR value is still given by (10), as long as a positive steady state exists. Note that Corollary 6.2 could be applied even if not all chemical reactions are known, and even if the model is not mass-action. It is also worth noting that the deficiency of the model is not known, due to the lack of information on the precise reactions constituting the network, but is certainly greater than 1. Indeed, the deficiency of the sub-system constituted by the EnvZ-OmpR osmoregulation system and by the misfolding of OmpR-P is 2, and the deficiency of a system is necessarily greater than or equal to the deficiency of any sub-system [15, Lemma 5].

## 7 Discussion

We have shown in Theorem 4.2 that a linear CI always exists for a family of ACR systems, and that this family strictly includes the models studied in [40]. The result, a more general version of which is proven in the Supplementary Material, has three main consequences: first, it provides an easy way to calculate the ACR value of ACR species. Secondly, as expressed in Theorem 6.1, the presence of a linear CI implies that the system is robust to arbitrary disturbances that do not vanish over time, under certain conditions. This fact can be naturally exploited to design perfect insulators, which are able to reject loading effects originated from the downstream modules. Finally, as expressed in Corollary 6.2, we are able to prove that, under certain conditions, the absolute concentration robustness of a portion of a system can be lifted to the whole model, and the ACR value of the ACR species remains unchanged.

The theory we developed opens the path for future research directions. First, efficient algorithms can be designed in order to check for the existence of portions of the systems that confer absolute concentration robustness to the whole system. To this aim, the connections of 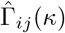 with structural properties of the network which we show in the Supplementary Material can be useful, and theoretical results can be expanded in this direction. Second, further detailed analysis on when stability can be ensured would be welcome. Currently, in the statement of Theorem 6.1 we cannot exclude the possibility that some species is completely consumed or indefinitely produced upon tampering with the model. Finding structural conditions able to eliminate this possibility would be a nice and useful contribution.

As a final remark, we think the study of stochastically modeled systems that satisfy the assumptions of Theorem 4.2 would be interesting and fruitful. Stochastic models of reaction systems are tipycally used when few molecules of certain chemical species are available [6, 20]. It is proven in [5] that systems satisfying the assumptions of Theorem 3.1, when stochastically modeled, undergo an extinction event almost surely. As a consequence, the desirable robustness properties of the ACR systems studied in [40] are completely destroyed in a low molecule copy-number regime. As an example, the model depicted in (1) undergoes an almost sure extinction of the chemical species *B* when stochastically modeled, regardless the initial conditions. This is caused by the fact that all the molecules of *B* can be consumed by the reaction *B → A*, before the occurrence of a reaction *A* + *B* → 2*B*. Robustness at finite time intervals of some stochastically modeled ACR systems is recovered, but only in a multiscale limit sense [4]. Moreover, it is shown in [3] that absolute concentration robustness of the deterministic model does not necessarily imply an extinction event in the corresponding stochastic model, but the connection is 13 largely unexplored. The results developed in the present paper can help in this direction: consider again (1). The extinction of species *B* cannot occur if production of *B* is included in the model as in (3), or as in

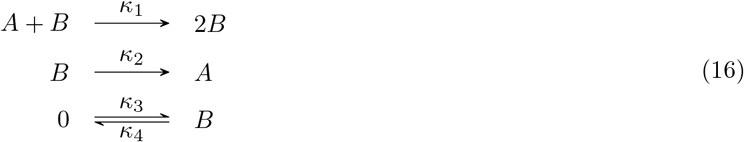

At the same time, it follows from Theorem 6.1 that the stability properties of the species *A* are maintained both in (3) and in (16), when deterministically modeled. In particular, the concentration of the species *A* still converges to the value *κ*_2_/*κ*_1_. It would be interesting to study if in this and in similar cases some form of absolute concentration robustness arise in the long-term dynamics of the stochastic models as well.

## Acknowledgement

This project has received funding from the European Research Council (ERC) under the European Union’s Horizon 2020 research and innovation programme grant agreement no. 743269 (CyberGenetics project).

## References

[1] U Alon, MG Surette, N Barkai, and S Leibler. Robustness in bacterial chemotaxis. Nature, 397(6715):168, 1999.

[2] DF Anderson. A modified next reaction method for simulating chemical systems with time dependent propensities and delays. J Chem Phys, 127(21):214107, 2007.

[3] DF Anderson and D Cappelletti. Discrepancies between extinction events and boundary equilibria in reaction networks. arXiv preprint arXiv:1809.04613, 2018.

[4] DF Anderson, D Cappelletti, and TG Kurtz. Finite time distributions of stochastically modeled chemical systems with absolute concentration robustness. SIAM J Appl Dyn Syst, 16(3):1309–1339, 2017.

[5] DF Anderson, GA Enciso, and MD Johnston. Stochastic analysis of biochemical reaction networks with absolute concentration robustness. J R Soc Interface, 11(93):20130943, 2014.

[6] DF Anderson and TG Kurtz. Stochastic analysis of biochemical systems, volume 1. Springer, 2015.

[7] D Angeli, JE Ferrell, and ED Sontag. Detection of multistability, bifurcations, and hysteresis in a large class of biological positive-feedback systems. Proc Natl Acad Sci U S A, 101(7):1822–1827, 2004.

[8] KJ Aström and RM Murray. Feedback systems: an introduction for scientists and engineers. Princeton university press, 2010.

[9] M Banaji and C Pantea. The inheritance of nondegenerate multistationarity in chemical reaction networks. SIAM J Appl Math, 78(2):1105–1130, 2018.

[10] N Barkai and S Leibler. Robustness in simple biochemical networks. Nature, 387(6636):913, 1997.

[11] E Batchelor and M Goulian. Robustness and the cycle of phosphorylation and dephosphorylation in a two-component regulatory system. Proc Natl Acad Sci U S A, 100(2):691–696, 2003.

[12] F Blanchini and E Franco. Structurally robust biological networks. BMC Syst Biol, 5(1):74, 2011.

[13] JD Brunner and G Craciun. Robust persistence and permanence of polynomial and power law dynamical systems. SIAM J Appl Math, 78(2):801–825, 2018.

[14] D Cappelletti, AP Majumder, and C Wiuf. Fixed–time and long–term dynamics of monomolecular reaction networks in stochastic environment. in preparation, 2019.

[15] D Cappelletti and C Wiuf. Product-form Poisson–like distributions and complex balanced reaction systems. SIAM J Appl Math, 76(1):411–432, 2016.

[16] G Craciun, F Nazarov, and C Pantea. Persistence and permanence of mass-action and power-law dynamical systems. SIAM J Appl Math, 73(1):305–329, 2013.

[17] D Del Vecchio, AJ Ninfa, and ED Sontag. Modular cell biology: retroactivity and insulation. Mol Syst Biol, 4(1):161, 2008.

[18] JP Dexter, T Dasgupta, and J Gunawardena. Invariants reveal multiple forms of robustness in bifunctional enzyme systems. Integrative Biology, 7(8):883–894, 2015.

[19] JC Doyle, BA Francis, and AR Tannenbaum. Feedback control theory. Courier Corporation, 2013.

[20] P Érdi and J Tóth. Mathematical models of chemical reactions: theory and applications of deterministic and stochastic models. Manchester University Press, 1989.

[21] M Feinberg. Chemical reaction network structure and the stability of complex isothermal reactors—i. the deficiency zero and deficiency one theorems. Chem Eng Sci, 42(10):2229–2268, 1987.

[22] M Feinberg. Foundations of Chemical Reaction Network Theory. Springer, 2019.

[23] M Feinberg and F Horn. Chemical mechanism structure and the coincidence of the stoichiometric and kinetic subspaces. Arch Ration Mech Anal, 66(1):83–97, 1977.

[24] E Feliu, D Cappelletti, and C Wiuf. Node balanced steady states: Unifying and generalizing complex and detailed balanced steady states. Math Biosci, 2018.

[25] E Feliu and C Wiuf. Enzyme–sharing as a cause of multi–stationarity in signalling systems. J R Soc Interface, page 20110664, 2011.

[26] S Feng, M Sáez, C Wiuf, E Feliu, and OS Soyer. Core signalling motif displaying multistability through multi-state enzymes. J R Soc Interface, 13(123):20160524, 2016.

[27] E Franco and F Blanchini. Structural properties of the MAPK pathway topologies in PC12 cells. J Math Biol, 67(6–7):1633–1668, 2013.

[28] M Gopalkrishnan, E Miller, and A Shiu. A geometric approach to the global attractor conjecture. SIAM J Appl Dyn Syst, 13(2):758–797, 2014.

[29] E Gross, HA Harrington, N Meshkat, and A Shiu. Joining and decomposing reaction networks. arXiv preprint arXiv:1810.05575, 2018.

[30] LH Hartwell, JJ Hopfield, S Leibler, and AW Murray. From molecular to modular cell biology. Nature, 402(6761supp):C47, 1999.

[31] T Jahnke and W Huisinga. Solving the chemical master equation for monomolecular reaction systems analytically. J Math Biol, 54(1):1–26, 2007.

[32] B Joshi and A Shiu. Atoms of multistationarity in chemical reaction networks. J Math Chem, 51(1):153–178, 2013.

[33] RL Karp, MP Millán, T Dasgupta, A Dickenstein, and J Gunawardena. Complex-linear invariants of biochemical networks. J Theor Biol, 311:130–138, 2012.

[34] D Mishra, PM Rivera, A Lin, D Del Vecchio, and R Weiss. A load driver device for engineering modularity in biological networks. Nat Biotechnol, 32(12):1268, 2014.

[35] T Miyashiro and M Goulian. High stimulus unmasks positive feedback in an autoregulated bacterial signaling circuit. Proc Natl Acad Sci U S A, 105(45):17457–17462, 2008.

[36] L Pantoja-Hernández and JC Martínez-García. Retroactivity in the context of modularly structured biomolecular systems. Front Bioeng Biotechnol, 3:85, 2015.

[37] LA Pratt and TJ Silhavy. Porin regulon of escherichia coli. In Two-Component Signal Transduction, pages 105–127. American Society of Microbiology, 1995.

[38] PEM Purnick and R Weiss. The second wave of synthetic biology: from modules to systems. Nat Rev Mol Cell Biol, 10(6):410, 2009.

[39] G Shinar, U Alon, and M Feinberg. Sensitivity and robustness in chemical reaction networks. SIAM J Appl Math, 69(4):977–998, 2009.

[40] G Shinar and M Feinberg. Structural sources of robustness in biochemical reaction networks. Science, 327(5971):1389–1391, 2010.

[41] G Shinar, R Milo, MR Martínez, and U Alon. Input–output robustness in simple bacterial signaling systems. Proc Natl Acad Sci U S A, 104(50):19931–19935, 2007.

[42] G Shinar, JD Rabinowitz, and U Alon. Robustness in glyoxylate bypass regulation. PLoS Comput Biol, 5(3):e1000297, 2009.

[43] AM Stock, VL Robinson, and PN Goudreau. Two-component signal transduction. Annu Rev Biochem, 69(1):183–215, 2000.

[44] K Takahashi, S Tănase-Nicola, and PR Ten Wolde. Spatio–temporal correlations can drastically change the response of a mapk pathway. Proc Natl Acad Sci U S A, 107(6):2473–2478, 2010.

[45] J Tóth, AL Nagy, and D Papp. Reaction kinetics: exercises, programs and theorems. Springer, 2018.

[46] F Xiao and JC Doyle. Robust perfect adaptation in biomolecular reaction networks. bioRxiv, page 299057, 2018.

[47] TM Yi, Y Huang, MI Simon, and JC Doyle. Robust perfect adaptation in bacterial chemotaxis through integral feedback control. Proc Natl Acad Sci U S A, 97(9):4649–4653, 2000.

