## Supplementary Material for "A hidden integral structure endows Absolute Concentration Robust systems with resilience to dynamical concentration disturbances"

### A Notation

#### A.1 General notation

We will denote by $\mathbb{R}$, $\mathbb{R}_{>0}$, and $\mathbb{R}_{\geq 0}$ the real, positive real, and non-negative real numbers, respectively. Similarly, we will denote by $\mathbb{Z}$, $\mathbb{Z}_{>0}$, and $\mathbb{Z}_{\geq 0}$ the integer, positive integer, and non-negative integer numbers, respectively. Given a real number $a$, we will denote its absolute value by $|a|$.

We denote by $e_i$ the vector that has 1 in its $i$th entry and 0 in all other entries. The dimension of such vector will be clear from the context, and if not it will be made explicit. Given two real vectors $v$, $w$ of the same length $n$, we denote their scalar product by $\langle v, w\rangle$, and we use the shorthand notation

$$v^w = \prod_{i=1}^n v_i^{w_i},$$

where $0^0$ is considered to be 1. We will denote the euclidean norm of $v$ by $\|v\|$.

Given two subsets $V, W \subseteq \mathbb{R}^n$, we denote their direct sum by

$$V \oplus W = \{v + w \, : \, v \in V, w \in W\}.$$

Moreover, given a vector $v \in \mathbb{R}^n$, we define

$$v + W = \{v + w \, : \, w \in W\}.$$

#### A.2 Reaction network terminology

##### A.2.1 Standard notation

A reaction network $\mathcal{G}$ is a triple $\{\mathcal{X}, \mathcal{C}, \mathcal{R}\}$, where

- $\mathcal{X}$ is a finite ordered set of $d$ different symbols, called *species*;
- $\mathcal{C}$ is a finite ordered set of $m$ linear combinations of species with non-negative integer coefficients, referred to as *complexes* and identified with vectors in $\mathbb{Z}^d_{\geq 0}$;
- $\mathcal{R}$ is a finite ordered set of elements of $\mathcal{C} \times \mathcal{C}$, referred to as *reactions*.

We will denote by $X_i$ the $i$th species of $\mathcal{X}$, and by $y_j$ the $j$th complex of $\mathcal{C}$, for $1 \leq i \leq d$ and $1 \leq j \leq m$. A reaction $(y_i, y_j)$ will be denoted by $y_i \to y_j$, and for any $y_i \in \mathcal{C}$ we assume there is no reaction of the form $y_i \to y_i$.

As mentioned, a complex $y_i$ can be regarded as a vector of $\mathbb{Z}^d_{\geq 0}$. Specifically, this is done by considering the $j$th entry $y_{ij}$ as the coefficient of $y_i$ relative to the $j$th species. In this regards, for any vector $v \in \mathbb{R}^d$ we will denote by $\operatorname{supp} v$ the subset of species such that

$$X_i \in \operatorname{supp} v \text{ if and only if } v_i \neq 0.$$

Similarly, for any vector $w \in \mathbb{R}^m$ we will denote by $\operatorname{supp} w$ the subset of complexes such that

$$y_i \in \operatorname{supp} v \text{ if and only if } v_i \neq 0.$$

We denote by

$$\mathcal{S} = \operatorname{span}_{\mathbb{R}}\{y_j - y_i \, : \, y_i \to y_j \in \mathcal{R}\}, \qquad \mathcal{S}^\perp = \{h \in \mathbb{R}^d \, : \, \langle h, y_j - y_i\rangle = 0 \text{ for all } y_i \to y_j \in \mathcal{R}\}.$$

The subspace $\mathcal{S}$ is called *stoichiometric subspace*, and the elements of $\mathcal{S}^\perp$ are called *conservation laws*.

A choice of kinetics for a reaction network is a set of (time-dependent) *rate functions* $\lambda_{ij} \colon \mathbb{R}^d_{\geq 0} \times \mathbb{R}_{\geq 0} \to \mathbb{R}_{\geq 0}$ for all $1 \leq i, j \leq m$, such that $\lambda_{ij}$ is a zero function if and only if $y_i \to y_j \notin \mathcal{R}$. A reaction network with a choice of kinetics $\mathscr{S} = (\mathcal{X}, \mathcal{C}, \mathcal{R}, \{\lambda_{ij}\}_{1 \leq i,j \leq m})$ is termed *reaction system.* A reaction system is associated with the system of differential equations

$$\frac{d}{dt}x(t) = \sum_{1 \leq i,j \leq m} (y_j - y_i)\lambda_{ij}(x(t), t) \quad \text{for all } t \in \mathbb{R}_{\geq 0}.$$

We note that rate functions are commonly intended to not depend on time, but we consider this more general setting in this paper.

A reaction system is called *mass-action system*, and denoted by $\mathscr{S} = (\mathcal{X}, \mathcal{C}, \mathcal{R}, \kappa)$, if there is a matrix $\kappa \in \mathbb{R}^{m\times m}_{\geq 0}$ such that

$$\lambda_{ij}(x,t) = \kappa_{ij} x^{y_i} \quad \text{for all } 1 \leq i,j \leq m, x \in \mathbb{R}^d_{\geq 0}.$$

In this case, the constants $\kappa_{ij}$ are termed *rate constants.* Define $\Lambda(x) \in \mathbb{R}^d_{\geq 0}$ by

$$\Lambda_i(x) = x^{y_i} \quad \text{for all } 1 \leq i \leq m, x \in \mathbb{R}^d_{\geq 0},$$

and define the $m \times m$ matrix $A(\kappa)$ by

$$A(\kappa)_{ij} = \begin{cases} \kappa_{ji} & \text{if } i \neq j \\ -\sum_{l=1}^m \kappa_{il} & \text{if } i = j \end{cases}$$

Finally, let $Y$ be the $d \times m$ matrix $Y$ with entries $Y_{ij} = y_{ji}$. Then, for mass-action sytems we have

$$\frac{d}{dt}x(t) = \sum_{1\leq i,j\leq m} (y_j - y_i)\kappa_{ij} x(t)^{y_i} = YA(\kappa)\Lambda(x(t)) \quad \text{for all } t \in \mathbb{R}_{\geq 0}.$$

The directed graph $\{\mathcal{C}, \mathcal{R}\}$ is called *reaction graph.* Reaction systems are often presented through the reaction graph, where indication on the reaction rates are written on top of the arrows that correspond to the related reaction. Specifically, the arrow corresponding to $y_i \to y_j$ is labeled with

$$\frac{\lambda_{ij}(x,t)}{x^y},$$

which corresponds to the rate constants for reaction rates of mass-action type. As an example, see (1) and (5) in the main text, or Figure 1. We denote by $\ell$ the number of connected components of the reaction graph. We define the *deficiency* as the number

$$\delta = m - \ell - \dim \mathcal{S}.$$

A *terminal component* of the reaction graph is a set of complexes $\mathcal{T} \subseteq \mathcal{C}$, such that

- if $y \in \mathcal{T}$ and there is a directed path in the reaction graph from $y$ to another complex $y'$, then $y' \in \mathcal{T}$;
- for any two complexes $y, y' \in \mathcal{T}$ with $y \neq y'$, there is a directed path from $y$ to $y'$, and a directed path from $y'$ to $y$.

Let $\tau$ be the number of terminal components, and denote by $\mathcal{T}^1, \mathcal{T}^2, \ldots, \mathcal{T}^\tau$ the different terminal components of the network. Since each connected component contains at least one terminal component, we have $\tau \geq \ell$. We say a complex is *terminal* if it is contained in a terminal component, and we say that a complex is *non-terminal* otherwise. As an example, the terminal components in (1) are $\{2B\}$ and $\{A\}$, hence the terminal complexes are $2B$ and $A$.

#### A.3 Absolute concentration robustness

**Definition A.1.** Consider a reaction system $\mathscr{S} = (\mathcal{X}, \mathcal{C}, \mathcal{R}, \{\lambda_{ij}\}_{1\leq i,j\leq m})$. A species $X_n \in \mathcal{X}$ is said to be *absolutely concentration robust* (ACR) if there exists $q \in \mathbb{R}_{>0}$ such that $c_n = q$ for all positive steady state $c$ of $\mathscr{S}$. In this case, $q$ is referred to as the *ACR value* of $X_n$. Finally, if an ACR species exists, then the reaction system $\mathscr{S}$ is said to be *ACR.*

*Remark* A.1. By definition, if a reaction system has no positive steady states, or a unique one, then all the species are ACR. We can call this a *degenerate case*, since, especially if no positive steady states exist, the concept of robustness is lost.

#### A.4 Control Theory terminology

Consider a differential equation of the form

$$\frac{d}{dt}x(t) = f(x(t)) + g(x(t), u(t)) \quad \text{for all } t \in \mathbb{R}_{\geq 0},$$

where $x\colon \mathbb{R}_{\geq 0} \to \mathbb{R}^d$ and $u\colon \mathbb{R}_{\geq 0} \to \mathbb{R}^{n_u}$ for some $n_u \in \mathbb{Z}_{>0}$, and $f$ and $g$ are differentiable functions. The function $u$ is called the *input of the system.* Further, we define *output of the system* the quantity $z(t) = a(x(t))$, for some differentiable function $a$, with $a\colon \mathbb{R}^d \to \mathbb{R}^{n_z}$ and $n_z \in \mathbb{Z}_{>0}$. Usually, one needs to find an appropriate function $u$ such that $z$ is close to a desired level $\overline{z} \in \mathbb{R}^{n_z}$, either on average or for $t \to \infty$. To this aim, the existence of a function $\phi\colon \mathbb{R}^{n_x} \to \mathbb{R}$ such that

$$\frac{d}{dt}\phi(x(t)) = z(t) - \overline{z}$$

is of high importance, an is called an *integral feedback* (IF) [8, 19]. If a function $\tilde{\phi}\colon \mathbb{R}^{n_x} \to \mathbb{R}$ satisfies

$$\frac{d}{dt}\tilde{\phi}(x(t)) = r(x(t))\Big(z(t) - \overline{z}\Big)$$

for some differentiable function $r\colon \mathbb{R}^{n_x} \to \mathbb{R}$, then $\tilde{\phi}$ is called a *constrained integral feedback* (CIF) [46].

In the setting of ACR species, $z(t)$ is usually the concentration of the ACR species at time $t$, and $\overline{z}$ is its ACR value.

### B Known results

The following result appears in the Appendix of [23].

**Theorem B.1.** *All the vectors in* $\ker A(\kappa)$ *have support in the terminal complexes. Specifically, there exists a basis* $\{\chi^1(\kappa), \chi^2(\kappa), \ldots, \chi^\tau(\kappa)\}$ *of* $\ker A(\kappa)$ *such that* $\operatorname{supp}\chi^i(\kappa) = \mathcal{T}^i$ *for all* $1 \leq i \leq m$.

The following result can be deduced from Section 6 in [23].

**Theorem B.2.** *We have*

$$\delta \geq \dim\Big(\ker Y \cap \operatorname{Im} A(\kappa)\Big) = \dim\ker YA(\kappa) - \dim\ker A(\kappa).$$

Finally, the following result is proven in the supplementary material of [40], where it is stated as Theorem S3.15.

**Theorem B.3.** *Consider a mass-action system* $(\mathcal{X}, \mathcal{C}, \mathcal{R}, \kappa)$*. Assume the following conditions hold:*

1. $y_i$ *and* $y_j$ *are non-terminal complexes;*
2. *the deficiency is 1;*

*Then, there exists* $q \in \mathbb{R}_{>0}$ *such that* $c^{y_j - y_i} = q$ *for all positive steady states* $c$*.*

For convenience, we restate here the weaker version given in the main text:

**Theorem 3.1.** *Consider a mass-action system, and assume the following holds:*

1. *there are two non-terminal complexes* $y_i$ *and* $y_j$ *such that only one entry of* $y_j - y_i$ *is non-zero;*
2. *the deficiency is 1.*
3. *a positive steady state exists.*

*Then, the species relative to the non-zero entry of* $y_j - y_i$ *is ACR.*

Note how Theorem 3.1 is a straightforward consequence of Theorem B.3: if the two complexes only differ in the $n$th entry, then

$$c^{y_j - y_i} = c_n^{(y_j - y_i)_n} = q$$

for all positive steady states $c$, hence $X_n$ is ACR.

*Remark* B.1. In the original statement of Theorem B.3, the additional hypothesis that a positive steady state exists is made. In fact, under the condition of Theorem B.3, the existence of positive steady states is not guaranteed. However, if no positive steady state exists then the conclusion of the theorem holds trivially. Hence, our reformulation holds correct. See Remark A.1 for the formal relationship between ACR species and the lack of positive steady states.

### C Calculation and structural properties of $\Gamma_{ij}(\kappa)$

Consider a mass-action system $\mathscr{S} = (\mathcal{X}, \mathcal{C}, \mathcal{R}, \kappa)$. For any $1 \leq i, j \leq m$ with $i \neq j$ define the set

$$\Gamma_{ij}(\kappa) = \left\{\gamma \in \mathbb{R}^{d+1} \,:\, \left(A(\kappa)^\top Y^\top \,|\, e_i\right)\gamma = e_j\right\}, \tag{C.1}$$

and for any real vector $v$ of length larger than $d$, let $\pi_d(v)$ be its projection onto the first $d$ components. We define

$$\hat{\Gamma}_{ij}(\kappa) = \left\{\hat{\gamma} \in \mathbb{R}^{d} \,:\, \hat{\gamma} = \pi_d(\gamma) \text{ for some } \gamma \in \Gamma_{ij}(\kappa)\right\}. \tag{C.2}$$

The set $\Gamma_{ij}(\kappa)$ can be computed by first calculating a basis for

$$\Psi_{ij}(\kappa) = \ker\left(A(\kappa)^\top Y^\top \,|\, e_i \,|\, e_j\right). \tag{C.3}$$

Note that this can be easily and quickly done by using a programming language which is able to deal with symbolic linear algebra, such as Matlab. The following holds

**Proposition C.1.** *Consider a mass-action system $\mathscr{S} = (\mathcal{X}, \mathcal{C}, \mathcal{R}, \kappa)$, and $1 \leq i, j \leq m$ with $i \neq j$. Let $\{\psi^1, \psi^2, \ldots, \psi^k\}$ be a basis for $\Psi_{ij}(\kappa)$, as defined in (C.3). Then, $\Gamma_{ij}(\kappa)$ (and consequently $\hat{\Gamma}_{ij}(\kappa)$) is non-empty if and only if $\psi^n_{d+2} \neq 0$ for some $1 \leq n \leq k$. If this is the case, then*

$$\Gamma_{ij}(\kappa) = \left\{-\frac{1}{\sum_{n=1}^k a_n\psi^n_{d+2}}\sum_{n=1}^k a_n\pi_{d+1}(\psi^n) \,:\, a_1, a_2, \ldots, a_k \in \mathbb{R} \text{ and } \sum_{n=1}^k a_n\psi^n_{d+2} \neq 0\right\},$$

*where $\pi_{d+1}\colon \mathbb{R}^{d+2} \to \mathbb{R}^{d+1}$ is the projection onto the first $d+1$ components, and*

$$\hat{\Gamma}_{ij}(\kappa) = \left\{-\frac{1}{\sum_{n=1}^k a_n\psi^n_{d+2}}\sum_{n=1}^k a_n\pi_{d}(\psi^n) \,:\, a_1, a_2, \ldots, a_k \in \mathbb{R} \text{ and } \sum_{n=1}^k a_n\psi^n_{d+2} \neq 0\right\}.$$

*Proof.* First, note that $\gamma \in \Gamma_{ij}(\kappa)$ if and only $(\gamma, -1) \in \Psi_{ij}(\kappa)$. This can be easily deduced by the definitions (C.1) and (C.3). The proof simply follows from this equivalence, and from the definition of $\hat{\Gamma}_{ij}(\kappa)$ given in (C.2). □

In Section F we will use Proposition C.1 to calculate $\hat{\Gamma}_{ij}(\kappa)$ for the main examples discussed in the main text. We will see that some vectors in the basis of $\Psi_{ij}(\kappa)$ do not depend on the particular choice of rate constants. As a consequence, some dynamical properties implied by the theory developed in this paper will only depend on the structure of the model rather than on a fine tuning of the parameters, which is desirable. In the following result, we explicitly derive structural properties of $\hat{\Gamma}_{ij}(\kappa)$.

**Proposition C.2.** *Consider a mass-action system $\mathscr{S} = (\mathcal{X}, \mathcal{C}, \mathcal{R}, \kappa)$, and $1 \leq i, j \leq m$ with $i \neq j$. Assume that $\hat{\Gamma}_{ij}(\kappa)$ is non-empty, and let $\hat{\gamma} \in \hat{\Gamma}_{ij}(\kappa)$. Then,*

$$\hat{\gamma} + \mathcal{S}^\perp \subseteq \hat{\gamma} + \ker A(\kappa)^\top Y^\top \subseteq \hat{\Gamma}_{ij}(\kappa). \tag{C.4}$$

*Moreover, if the mass-action system has a steady state $c$ with $c^{y_i} > 0$, then*

$$\gamma'_{d+1} = c^{y_j - y_i} \quad \text{for all } \gamma' \in \Gamma_{ij}(\kappa) \tag{C.5}$$

*and*

$$\hat{\Gamma}_{ij}(\kappa) = \hat{\gamma} + \ker A(\kappa)^\top Y^\top. \tag{C.6}$$

*Finally, if each connected component of the reaction graph contains exactly one terminal component, then*

$$\ker A(\kappa)^\top Y^\top = \mathcal{S}^\perp. \tag{C.7}$$

*Proof.* First, we have that for any $\hat{\gamma} \in \hat{\Gamma}_{ij}(\kappa)$

$$\hat{\gamma} + \ker A(\kappa)^\top Y^\top \subseteq \hat{\Gamma}_{ij}(\kappa). \tag{C.8}$$

Indeed, for any $\hat{\gamma} \in \hat{\Gamma}_{ij}(\kappa)$, there exists $\gamma \in \Gamma_{ij}(\kappa)$ with $\hat{\gamma} = \pi_d(\gamma)$. Then, for any for any $v \in \ker A(\kappa)^\top Y^\top$ we have

$$\left(A(\kappa)^\top Y^\top \,|\, e_i\right)\left(\gamma + \begin{pmatrix} v \\ 0 \end{pmatrix}\right) = \left(A(\kappa)^\top Y^\top \,|\, e_i\right)\gamma + \begin{pmatrix} A(\kappa)^\top Y^\top v \\ 0 \end{pmatrix} = e_j.$$

Hence,

$$\gamma + \begin{pmatrix} v \\ 0 \end{pmatrix} \in \Gamma_{ij}(\kappa)$$

which implies that $\hat{\gamma} + v \in \hat{\Gamma}_{ij}(\kappa)$ and proves (C.8). Since for any $h \in \mathcal{S}^\perp$ and any $1 \le n \le m$

$$\left(A(\kappa)^\top Y^\top h\right)_n = \sum_{l=1}^m \langle h, y_l - y_n \rangle \kappa_{nl} = 0,$$

it follows

$$\mathcal{S}^\perp \subseteq \ker A(\kappa)^\top Y^\top. \tag{C.9}$$

(C.4) follows from (C.8) and (C.9).

Now assume that the mass-action system has a steady state $c$, with $c^{y_i} > 0$. Then,

$$0 = \frac{d}{dt}\langle \hat{\gamma}, x(t) \rangle|_{x(t)=c} = \hat{\gamma}^\top Y A(\kappa)\Lambda(c) = (e_j - \gamma_{d+1} e_i)\Lambda(c) = c^{y_j} - \gamma_{d+1} c^{y_i}.$$

Since $c^{y_i} > 0$, then necessarily $\gamma_{d+1} = c^{y_j - y_i}$. For the same argument, for any other $\gamma' \in \Gamma_{ij}(\kappa)$, $\gamma'_{d+1} = \gamma_{d+1} = c^{y_j - y_i}$. Hence, (C.5) holds. It follows that any $\gamma' \in \Gamma_{ij}(\kappa)$ is of the form

$$\gamma' = \gamma + \begin{pmatrix} v \\ 0 \end{pmatrix}$$

for some $v \in \mathbb{R}^d$. Moreover, since $\gamma' \in \Gamma_{ij}(\kappa)$

$$e_j = \left(A(\kappa)^\top Y^\top \,|\, e_i\right)\gamma' = \left(A(\kappa)^\top Y^\top \,|\, e_i\right)\gamma + \begin{pmatrix} A(\kappa)^\top Y^\top v \\ 0 \end{pmatrix} = e_j + \begin{pmatrix} A(\kappa)^\top Y^\top v \\ 0 \end{pmatrix}.$$

Hence, necessarily $v \in \ker A(\kappa)^\top Y^\top$, which implies

$$\hat{\Gamma}_{ij}(\kappa) \subseteq \hat{\gamma} + \ker A(\kappa)^\top Y^\top.$$

The latter, together with (C.8), implies (C.6).

To conclude the proof, we need to show that if each connected component of the reaction graph contains exactly one terminal component (i.e. if $\ell = \tau$), then (C.7) holds. This follows from (C.9) and

$$\begin{aligned} \dim \mathcal{S}^\perp &= d - \dim \mathcal{S} = d + \ell + \delta - m \\ &= d + \tau + \delta - m \ge d - m + \dim\ker Y A(\kappa) \\ &= \dim\ker A(\kappa)^\top Y^\top, \end{aligned}$$

where we utilized Theorems B.1 and B.2 for the forth equality. □

As a consequence, we have the following.

**Corollary C.3.** *Consider a mass-action system, and assume that $y_i$ and $y_j$ are two complexes for which $\hat{\Gamma}_{ij}(\kappa)$ is non-empty. Let $\{v_p\}_{p=1}^H$ be a basis for $\mathcal{S}^\perp$, where $H = d - \dim \mathcal{S}$. Moreover, let $X_{l_1}, X_{l_2}, \dots, X_{l_n} \in \mathcal{X}$ such that the rank $H \times n$ matrix $V$ has rank $n$, where $V_{pq} = v_{pl_q}$ for all $1 \le p \le H$ and $1 \le q \le n$. Then, there exists $\hat{\gamma} \in \hat{\Gamma}_{ij}(\kappa)$ such that $\langle \hat{\gamma}, e_p \rangle = 0$ for all $1 \le p \le n$.*

*Proof.* Let $\hat{\gamma}^\star \in \hat{\Gamma}_{ij}(\kappa)$. It follows from Proposition C.2 that for all $w \in \mathbb{R}^H$

$$\hat{\gamma}^\star + \sum_{p=1}^H w_p v_p \in \hat{\Gamma}_{ij}(\kappa).$$

Hence, the proof is concluded by choosing $w$ such that

$$w^\top V = -(\hat{\gamma}^\star_{l_1}, \hat{\gamma}^\star_{l_2}, \dots, \hat{\gamma}^\star_{l_n}),$$

which is possible because the $H \times n$ matrix $V$ has rank $n$. □

Further useful structural property of $\hat{\Gamma}_{ij}(\kappa)$ are following. Such properties are useful while checking whether the results of this paper can be applied, and can be used by an algorithm designed to this aim. Before stating the structural results, it is convenient to prove the following lemma.

**Lemma C.4.** *Consider a mass-action system $\mathscr{S} = (\mathcal{X}, \mathcal{C}, \mathcal{R}, \kappa)$, and $1 \leq i, j \leq m$ with $i \neq j$. Then, $\hat{\Gamma}_{ij}(\kappa)$ is non-empty if and only if $e_j - \gamma_{d+1} e_i \in (\ker Y A(\kappa))^\top$ for some $\gamma_{d+1} \in \mathbb{R}$. Moreover, if that is the case and if the mass-action system has a steady state $c$ with $c^{y_i} > 0$ and $c^{y_j} > 0$, then necessarily $\gamma_{d+1} > 0$.*

*Proof.* By definition of $\hat{\Gamma}_{ij}(\kappa)$ given in (C.2), $\hat{\Gamma}_{ij}(\kappa)$ is non-empty if and only if $\Gamma_{ij}(\kappa)$ is non-empty. By (C.1), $\Gamma_{ij}(\kappa)$ is non-empty if and only if there exists $\gamma_{d+1} \in \mathbb{R}$ such that $e_j - \gamma_{d+1} e_i$ is in $\operatorname{Im} A(\kappa)^\top Y^\top$. By the fundamental theorem of linear algebra, the latter holds if and only if $e_j - \gamma_{d+1} e_i$ is orthogonal to $\ker Y A(\kappa)$.

To conclude the proof, we note that if the mass-action system has a steady state $c$ with $c^{y_i} > 0$ and $\hat{\Gamma}_{ij}(\kappa)$ is non-empty, then it follows by (C.5) in Proposition C.2 that the quantity $\gamma_{d+1}$ discussed in the first part of the proof is necessarily equal to $c^{y_j - y_i} > 0$. □

**Proposition C.5.** *Consider a mass-action system $\mathscr{S} = (\mathcal{X}, \mathcal{C}, \mathcal{R}, \kappa)$, and $1 \leq i, j \leq m$. Assume that one of the following holds:*

- *$y_i$ is non-terminal and $y_j$ is terminal;*
- *$y_i$ and $y_j$ are in two different terminal components.*

*Then, $\hat{\Gamma}_{ij}(\kappa)$ is empty.*

*Proof.* By assumption, $y_j$ is terminal. By Theorem B.1, there exists a vector

$$\chi \in \ker A(\kappa) \subseteq \ker Y A(\kappa)$$

such that $\operatorname{supp} \chi$ is the terminal component containing $y_j$. Hence, if $y_i$ is non-terminal or if $y_i$ is in a different terminal component than $y_j$, we have that for all $\gamma_{d+1} \in \mathbb{R}$

$$\langle \chi, e_j - \gamma_{d+1} e_i \rangle = \chi_j \neq 0.$$

It follows from Lemma C.4 that $\hat{\Gamma}_{ij}(\kappa)$ is empty, which concludes the proof. □

**Proposition C.6.** *Consider a mass-action system $\mathscr{S} = (\mathcal{X}, \mathcal{C}, \mathcal{R}, \kappa)$, and $1 \leq i, j \leq m$ with $i \neq j$. Assume that a steady state $c$ with $c^{y_i} > 0$ and $c^{y_j} > 0$ exists. Then, $\hat{\Gamma}_{ij}(\kappa)$ is empty if and only if $\hat{\Gamma}_{ji}(\kappa)$ is empty.*

*Proof.* By the symmetric role of $i$ and $j$, it suffices to prove that if $\hat{\Gamma}_{ij}(\kappa)$ is non-empty, then necessarily $\hat{\Gamma}_{ij}(\kappa)$ is non-empty.

By Lemma C.4, if $\hat{\Gamma}_{ij}(\kappa)$ is non-empty then there exists $\gamma_{d+1} \in \mathbb{R}$ with $e_j - \gamma_{d+1} e_i \in (\ker Y A(\kappa))^\top$, and $\gamma_{d+1} > 0$. Hence,

$$e_i - \frac{1}{\gamma_{d+1}} e_j \in (\ker Y A(\kappa))^\top,$$

hence by Lemma C.4 $\hat{\Gamma}_{ji}(\kappa)$ is non-empty, which is what we wanted to show. □

### D Proofs of the results stated in the main text

In this section, we state and prove more general versions of the theorems presented in the main text.

#### D.1 Existence and characterization of a CIF

The following result is stated in the main text.

**Theorem 4.1.** *Consider a mass-action system, and assume the following holds:*

1. *there are two non-terminal complexes $y_i$ and $y_j$ such that only one entry of $y_j - y_i$ is non-zero;*
2. *the deficiency is 1.*
3. *a positive steady state exists.*

*Then, $\hat{\Gamma}_{ij}(\kappa)$ is non-empty.*

The following more general result holds, from which Theorem 4.1 can be immediately deduced.

**Theorem D.1.** *Consider a mass-action system, and assume the following holds:*

1. *$y_i$ and $y_j$ are two distinct non-terminal complex;*

2. *the deficiency is 1;*

3. *a steady state $c$ with $c^{y_i} \neq 0$ exists.*

*Then, $\hat{\Gamma}_{ij}(\kappa)$ is non-empty.*

*Proof.* By condition 3, $c$ is a steady state, hence the vector $\Lambda(c)$ is in $\ker YA(\kappa)$. Moreover, $\Lambda_i(c) = c^{y_i} \neq 0$. By condition 1, $y_i$ is a non-terminal complex. Hence, by combining $\Lambda_i(c) \neq 0$ with Theorem B.1, it follows that $\Lambda(c)$ is not in $\ker A(\kappa)$. Since by assumption $\delta = 1$, it follows from Theorem B.2 that

$$\ker YA(\kappa) = \operatorname{span}_{\mathbb{R}}\{\Lambda(c)\} \oplus \ker A(\kappa). \tag{D.1}$$

Define

$$v = e_j - c^{y_j - y_i} e_i = e_j - \frac{\Lambda_j(c)}{\Lambda_i(c)} e_i.$$

Clearly, $v$ is orthogonal to $\Lambda(c)$. Moreover, by condition 1, both $y_i$ and $y_j$ are non-terminal complexes. Hence, the vector $v$ is orthogonal to $\ker A(\kappa)$ by Theorem B.1. It follows from (D.1) that $v$ is orthogonal to $\ker YA(\kappa)$, which implies that $\hat{\Gamma}_{ij}(\kappa)$ is non-empty by Lemma C.4. □

We now proceed to prove the following result, stated in the main text.

**Theorem 4.2.** *Consider a mass-action system. Assume that there are two complexes $y_i$ and $y_j$ only differing in the $n$th entry, and that $\hat{\Gamma}_{ij}(\kappa)$ is non-empty. Let $\gamma \in \hat{\Gamma}_{ij}(\kappa)$, and define*

$$q = \gamma_{d+1}^{\frac{1}{(y_j - y_i)_n}}.$$

*Then, either no positive steady state exists or the $n$th species is ACR with ACR value $q$. Moreover,*

$$\phi(x) = \sum_{i=1}^{d} \beta_i x_i$$

*is a linear CI with*

$$\frac{d}{dt}\phi(x(t)) = \Lambda_i(x(t)) \left( x_n(t)^{(y_j - y_i)_n} - q^{(y_j - y_i)_n} \right)$$

*for any initial condition $x(0)$ if and only if $\beta \in \hat{\Gamma}_{ij}(\kappa)$.*

In order to prove the result, we will show that a more general version holds, which we state here.

**Theorem D.2.** *Consider a mass-action system, and assume that $y_i$ and $y_j$ are two complexes for which $\hat{\Gamma}_{ij}(\kappa)$ is non-empty. Let $\gamma \in \hat{\Gamma}_{ij}(\kappa)$. Then,*

$$c^{y_j - y_i} = \gamma_{d+1} \tag{D.2}$$

*for all steady states $c$ satisfying $c^{y_i} \neq 0$. Moreover,*

$$\phi(x) = \sum_{i=1}^{d} \beta_i x_i$$

*is a linear CIF with*

$$\frac{d}{dt}\phi(x(t)) = \Lambda_i(x(t)) \Big( x(t)^{y_j - y_i} - \gamma_{d+1} \Big) \tag{D.3}$$

*for any initial condition $x(0)$ if and only if $\beta \in \hat{\Gamma}_{ij}(\kappa)$.*

*Proof.* (D.2) follows from (C.5) in Proposition C.2.

Let $\beta \in \hat{\Gamma}_{ij}(\kappa)$. Then, by Proposition C.2 we have that $\beta - \pi_d(\gamma) \in \ker A(\kappa)^\top Y^\top$. Hence, for any initial condition $x(0) \in \mathbb{R}^d_{\geq 0}$ and any $t \in \mathbb{R}_{\geq 0}$,

$$\frac{d}{dt}\langle \beta, x(t) \rangle = \beta^\top YA(\kappa)\Lambda(x(t)) = \pi_d(\gamma)^\top YA(\kappa)\Lambda(x(t)) = (e_j - \gamma_{d+1} e_i)^\top \Lambda(x(t)) = x(t)^{y_i} \Big( x(t)^{y_j - y_i} - \gamma_{d+1} \Big),$$

which is (D.3).

Conversely, assume that (D.3) holds. Then, for all $x \in \mathbb{R}^d_{\geq 0}$ we have

$$(\beta^\top - \pi_d(\gamma))^\top YA(\kappa)\Lambda(x) = 0.$$

Since the entries of $\Lambda$ are linearly independent monomials on $\mathbb{R}^d$, it must be $(\beta^\top - \pi_d(\gamma)) \in \ker A(\kappa)^\top Y^\top$, which implies that $\beta \in \hat{\Gamma}_{ij}(\kappa)$ by Proposition C.2. □

#### D.2 Rejection of persistent disturbances

We recall here the formal definition of "oscillation", as intended in this paper.

**Definition D.1.** We say that a function $g\colon \mathbb{R}_{\geq 0} \to \mathbb{R}$ *oscillates* around a value $\overline{q} \in \mathbb{R}$ if for each $t \in \mathbb{R}_{\geq 0}$ there exist $t_+ > t$ and $t_- > t$ such that

$$g(t_+) > q \quad \text{and} \quad g(t_-) < q.$$

The following result holds.

**Theorem D.3.** *Consider a mass-action system, with associated differential equation*

$$\frac{d}{dt}x(t) = f(x(t)).$$

*Assume that $y_i$ and $y_j$ are two complexes for which $\hat{\Gamma}_{ij}(\kappa)$ is non-empty. Then, there exists $\overline{q} \in \mathbb{R}_{\geq 0}$ such that $c^{y_j - y_i} = \overline{q}$ for all steady states $c$ with $c^{y_i} \neq 0$. Consider an arbitrary function $u\colon \mathbb{R}^d_{\geq 0} \times \mathbb{R}_{\geq 0} \to \mathbb{R}^d$ such that a solution to*

$$\frac{d}{dt}\tilde{x}(t) = f(\tilde{x}(t)) + u(\tilde{x}(t), t)$$

*exists with $\tilde{x}(t) \geq 0$ for all $t \geq 0$. Assume that there exists a $\hat{\gamma} \in \hat{\Gamma}_{ij}(\kappa)$ which is orthogonal to the vector $u(x,t)$ for any $x \in \mathbb{R}^d_{\geq 0}$, $t \in \mathbb{R}_{\geq 0}$. Then, for any initial condition $\tilde{x}(0) \in \mathbb{R}^d_{\geq 0}$, at least one of the following statements holds:*

1. *There is a species $X_k \in \operatorname{supp} \hat{\gamma}$ such that $\limsup_{t\to\infty} \tilde{x}_k(t) = \infty$;*
2. *There is a species $X_k \in \operatorname{supp} y_i$ such that $\liminf_{t\to\infty} \tilde{x}_k(t) = 0$;*
3. *$\tilde{x}(t)^{y_j - y_i}$ oscillates around $\overline{q}$;*
4. $\displaystyle\lim_{t\to\infty} \int_t^\infty \left|\tilde{x}(s)^{y_j - y_i} - \overline{q}\right| ds = 0.$

*Proof.* It follows from Theorem D.2 that there exists $\overline{q} \in \mathbb{R}_{\geq 0}$ such that $c^{y_j - y_i} = \overline{q}$ for all steady states $c$ with $c^{y_i} \neq 0$. Note that in this setting $c^{y_i} \neq 0$ is equivalent to $c^{y_i} > 0$, since the state space is limited to vectors of non-negative concentrations.

Fix an initial condition $\tilde{x}(0) \in \mathbb{R}^d_{\geq 0}$. Assume that 1 does not occur. Hence, there exists $M \in \mathbb{R}_{>0}$ such that

$$|\langle \hat{\gamma}, \tilde{x}(t) \rangle| < M \quad \text{for all } t \in \mathbb{R}_{\geq 0}.$$

By the fact that $\hat{\gamma}$ is orthogonal to $u(x,t)$ for all $x \in \mathbb{R}^d_{\geq 0}$ and all $t \geq 0$, and by Theorem D.2, we have

$$\frac{d}{dt}\langle \hat{\gamma}, \tilde{x}(t) \rangle = \langle \hat{\gamma}, f(\tilde{x}(t)) \rangle = \tilde{x}(t)^{y_i}\left(\tilde{x}(t)^{y_j - y_i} - \overline{q}\right).$$

It follows that

$$\left|\int_0^t \tilde{x}(s)^{y_i}\left(\tilde{x}(s)^{y_j - y_i} - \overline{q}\right) ds\right| < M - |\langle \hat{\gamma}, \tilde{x}(0) \rangle| \quad \text{for all } t \in \mathbb{R}_{\geq 0}. \tag{D.4}$$

If 2 does not hold, then there exists $m \in \mathbb{R}_{>0}$ such that $x(t)^{y_i} > m$ for all $t \in \mathbb{R}_{>0}$. Together with (D.4), this would imply that there exists $M^\star \in \mathbb{R}_{>0}$ such that

$$\left|\int_0^t \left(\tilde{x}(s)^{y_j - y_i} - \overline{q}\right) ds\right| < M^\star \quad \text{for all } t \in \mathbb{R}_{\geq 0}. \tag{D.5}$$

If 3 does not hold, then there exists $t^\star \in \mathbb{R}_{>0}$ such that $\tilde{x}(t)^{y_j - y_i} - \overline{q}$ maintains the same sign for all $t > t^\star$, which together with (D.5) implies 4. The proof is then concluded. □

Now we prove Theorem 6.1, which is stated in the main text and which we state here again for convenience. Theorem 6.1 follows almost entirely from Theorem D.3.

**Theorem 6.1.** *Consider a mass-action system, with associated differential equation*

$$\frac{d}{dt}x(t) = f(x(t)).$$

*Assume that there are two complexes $y_i$ and $y_j$ only differing in the nth entry, and that $\hat{\Gamma}_{ij}(\kappa)$ is non-empty. Let $q$ be the ACR value of the nth species. Consider an arbitrary function $u$ with image in $\mathbb{R}^d$ such that a solution to*

$$\frac{d}{dt}\tilde{x}(t) = f(\tilde{x}(t)) + u(\tilde{x}(t), t)$$

*exists. Assume that there exists a $\hat{\gamma} \in \hat{\Gamma}_{ij}(\kappa)$ which is orthogonal to the vector $u(x,t)$ for any $x, t$. Then, for any initial condition $\tilde{x}(0)$, at least one of the following holds:*

(a) *the concentration of some species goes to 0 or infinity, along a sequence of times;*

(b) $\tilde{x}_n(t)$ *oscillates around* $q$ *and* $\hat{\gamma}_k \neq 0$ *for some* $k \neq n$*;*

(c) *the integral*

$$\int_t^\infty |\tilde{x}_n(s) - q| ds$$

*tends to* $0$*, as* $t$ *goes to infinity.*

*Proof.* Assume (a) does not hold. Then, there exists $\varepsilon \in \mathbb{R}_{>0}$ such that

$$\varepsilon \leq \|\tilde{x}(t)\| \leq \frac{1}{\varepsilon} \quad \text{for all } t \in \mathbb{R}_{\geq 0}. \tag{D.6}$$

Moreover, it follows from Theorem D.3 that at least one of the following holds:

1. $\tilde{x}_n(t)^{(y_j - y_i)_n}$ oscillates around $q^{(y_j - y_i)_n}$, implying that $\tilde{x}_n(t)$ oscillates around $q$;

2. $\lim_{t\to\infty} \int_t^\infty \left|\tilde{x}_n(s)^{(y_j - y_i)_n} - q^{(y_j-y_i)_n}\right| ds = 0.$

Since 2 clearly implies (c), to complete the proof it suffices to show that 1 implies $\hat{\gamma} \neq \hat{\gamma}_n e_n$.

Assume 1 holds. If it were $\hat{\gamma} = \hat{\gamma}_n e_n$, then by Theorem D.2 and by the fact that $u(x,t)$ is orthogonal to $\hat{\gamma}$ for all $x \in \mathbb{R}^d_{\geq 0}$ we would have

$$\hat{\gamma}_n \frac{d}{dt} \tilde{x}_n(t) = \frac{d}{dt} \langle \hat{\gamma}, \tilde{x}(t) \rangle = \langle \hat{\gamma}, f(\tilde{x}(t)) \rangle = \tilde{x}(t)^{y_i} \Big( \tilde{x}_n(t) - q \Big) \quad \text{for all } t \in \mathbb{R}_{\geq 0}.$$

By (D.6), there would be $m, M \in \mathbb{R}_{>0}$ such that

$$m \Big( \tilde{x}_n(t) - q \Big) \leq \hat{\gamma}_n \frac{d}{dt} \Big( \tilde{x}_n(t) - q \Big) \leq M \Big( \tilde{x}_n(t) - q \Big) \quad \text{for all } t \in \mathbb{R}_{\geq 0},$$

which would in turn imply that $\tilde{x}_n(t) - q$ maintains the same sign for all $t \in \mathbb{R}_{\geq 0}$. Hence, 1 could not hold and the proof is concluded. □

#### D.3 Inclusion in larger systems

Loosely speaking, a subsystem $\mathscr{S}'$ of a reaction system $\mathscr{S}$ is simply the reaction system generated by a subset of the reactions of $\mathscr{S}$, with the same choice of rate functions. In order to give a formal definition, some more care is needed as the number of species involved in the subsystem may be smaller (hence the dimensions of $\mathscr{S}$ and $\mathscr{S}'$ may be different).

**Definition D.2.** Let $\mathscr{S} = (\mathcal{X}, \mathcal{C}, \mathcal{R}, \{\lambda_{ij}\}_{1 \leq i,j \leq m})$ be a reaction system with $d$ species and $m$ complexes. A *subsystem* of $\mathscr{S}$ is a reaction system $\mathscr{S}' = (\mathcal{X}', \mathcal{C}', \mathcal{R}', \{\lambda'_{ij}\}_{1 \leq i,j \leq m'})$ with $d'$ species and $m'$ complexes such that, up to reordering of $\mathcal{X}$ and $\mathcal{C}$,

- $\mathcal{X}' = \{X_i \in \mathcal{X} \,:\, 1 \leq i \leq d'\}$;
- $y_{in} = 0$ for all $y_i \in \mathcal{C}$ with $1 \leq i \leq m'$ and for all $d' < n \leq d$;
- $\mathcal{C}' = \{y'_i \,:\, y'_i = \pi(y_i), y_i \in \mathcal{C}, 1 \leq i \leq m'\}$, where $\pi : \mathbb{R}^d \to \mathbb{R}^{d'}$ is the projection onto the first $d'$ components;
- $\mathcal{R}' \subseteq \{y'_i \to y'_j \,:\, y_i \to y_j \in \mathcal{R}\}$;
- for all $1 \leq i,j \leq m'$ and all $x \in \mathbb{R}^d_{\geq 0}$

$$\lambda'_{ij}(\pi(x)) = \begin{cases} \lambda_{ij}(x) & \text{if } y'_i \to y'_j \in \mathcal{R}' \\ 0 & \text{otherwise.} \end{cases}$$

With a slight abuse of notation due to the potential different lenght of the complexes of $\mathscr{S}$ and $\mathscr{S}'$, we further say that $y_k \to y_l$ is a reaction of $\mathscr{S}$ but is not a reaction of $\mathscr{S}'$ if $y_k \to y_j \in \mathcal{R}$ and either $\max k, l > m'$ or $y'_k \to y'_l \notin \mathcal{R}'$. In this case, we write $y_k \to y_l \in \mathcal{R} \setminus \mathcal{R}'$.

The following result is stated in the main text.

**Corollary 6.2.** *Consider a reaction system $\mathscr{S}$, and let $\mathscr{S}'$ be a sub-system. Assume that $\mathscr{S}'$ is a mass-action system with two complexes $\pi(y_i)$ and $\pi(y_j)$ only differing in the entry relative to the species $X$, and for which $\hat{\Gamma}_{ij}(\kappa)$ is non-empty. Moreover, assume there exists $\hat{\gamma} \in \hat{\Gamma}_{ij}(\kappa)$ such that $\pi(y_l - y_k)$ is orthogonal to $\hat{\gamma}$ for all $y_k \to y_l$ that are reactions of $\mathscr{S}$ but not reactions of $\mathscr{S}'$. Then, the $X$ is ACR for both $\mathscr{S}$ and $\mathscr{S}'$, with the same ACR value.*

Here, we prove the following, stronger version.

**Corollary D.4.** *Consider a reaction system $\mathscr{S} = (\mathcal{X}, \mathcal{C}, \mathcal{R}, \{\lambda_{ij}\}_{1\leq i,j\leq m})$, and let $\mathscr{S}' = (\mathcal{X}', \mathcal{C}', \mathcal{R}', \{\lambda'_{ij}\}_{1\leq i,j\leq m'})$ be a sub-system. Assume that $\mathscr{S}'$ is a mass-action system with two complexes $\pi(y_i)$ and $\pi(y_j)$ for which $\hat{\Gamma}_{ij}(\kappa)$ is non-empty. Moreover, assume there exists $\hat{\gamma} \in \hat{\Gamma}_{ij}(\kappa)$ such that $\pi(y_l - y_k)$ is orthogonal to $\hat{\gamma}$ for all $y_k \to y_l \in \mathcal{R}\setminus\mathcal{R}'$. Then, there exists a value $\overline{q} \in \mathbb{R}_{\geq 0}$ such that*

$$(c')^{\pi(y_j - y_i)} = \overline{q}$$

*for all steady states $c'$ of $\mathscr{S}'$ such that $(c')^{\pi(y_i)} > 0$, and*

$$c^{y_j - y_i} = \overline{q}$$

*for all steady states $c$ of $\mathscr{S}$ such that $c^{y_i} > 0$.*

*Proof.* The existence of $\overline{q} \in \mathbb{R}_{\geq 0}$ such that

$$(c')^{\pi(y_j - y_i)} = \overline{q}$$

for all steady states $c'$ of $\mathscr{S}'$ with $(c')^{\pi(y_i)} > 0$ follows from Theorem D.2.

In order to prove the second part of the statement, we can write

$$\frac{d}{dt}\pi(x(t)) = f(\pi(x(t))) + u(x(t), t),$$

where

$$\begin{aligned} f(\pi(x(t))) &= \sum_{y'_k \to y'_l \in \mathcal{R}'} (y'_l - y'_k)\kappa_{ij}\pi(x(t))^{y_i} \\ u(x(t), t) &= \sum_{y_k \to y_l \in \mathcal{R}\setminus\mathcal{R}'} \pi(y_l - y_k)\lambda_{ij}(x(t), t). \end{aligned}$$

Note that $\hat{\gamma}$ is orthogonal to $u(x, t)$, for all $x \in \mathbb{R}^d_{\geq 0}$ and $t \in \mathbb{R}_{\geq 0}$. Hence, if $c$ is a steady state of $\mathscr{S}$, it follows from Theorem D.2 that

$$\begin{aligned} 0 &= \frac{d}{dt}\langle\hat{\gamma}, \pi(c)\rangle = \langle\hat{\gamma}, f(\pi(c))\rangle \\ &= \pi(c)^{\pi(y_i)}\Big(\pi(c)^{\pi(y_j - y_i)} - \overline{q}\Big) \\ &= c^{y_i}\Big(c^{y_j - y_i} - \overline{q}\Big), \end{aligned}$$

which concludes the proof. □

### E Example of an ACR system with no linear CIF

Assume that $X_i$ is an ACR species, with ACR value $q$. It is not always possible to find a *linear* combination of species whose derivative at time $t$ is of the form $r(t)(x_i(t)^\alpha - q)$, for some polynomial $r(t)$ and some real number $\alpha$. As an example, consider the following mass-action system:

$$\begin{array}{lclcl} \mathrm{A{+}B} & \xrightarrow{\kappa_1} & \mathrm{B{+}C} & \underset{\kappa_3}{\overset{\kappa_2}{\rightleftharpoons}} & \mathrm{2B} \\ \mathrm{B} & \underset{\kappa_5}{\overset{\kappa_4}{\rightleftharpoons}} & \mathrm{2E} & \xrightarrow{\kappa_6} & \mathrm{2D} \\ \mathrm{C} & \xrightarrow{\kappa_7} & \mathrm{A} & & \\ \mathrm{D} & \xrightarrow{\kappa_8} & \mathrm{E} & & \end{array}$$

Let us order the species alphabetically, and the complexes as $(A + B, B + C, 2B, B, 2E, 2D, C, A, D, E)$. It is proven in [3] that the species $A$ is ACR, with ACR value

$$q = \frac{\kappa_3\kappa_7}{\kappa_1\kappa_2}.$$

In this case there is no linear combination of species whose derivative is of the form

$$r(t)\left(x_1(t)^\alpha - q^\alpha\right), \tag{E.1}$$

for some polynomial function $r(t)$ and some real number $\alpha$. Indeed, for any $\beta \in \mathbb{R}^d$, the derivative of $\langle\beta, x(t)\rangle$ is given by $\beta^\top YA(\kappa)\Lambda(x(t))$, which in this case is a polynomial of the form

$$w_1x_1(t)x_2(t) + w_2x_3(t) + w_3x_2(t)x_3(t) + w_4x_2(t)^2 + w_5x_2(t) + w_6x_5(t)^2 + w_7x_4(t),$$

for some real coefficients $w_i$. The only monomial containing $x_1(t)$ is the first one, so if we want the derivative of $\langle\beta, x(t)\rangle$ to be of the form (E.1), necessarily $w_1 \neq 0$, $\alpha = 1$, and $r(t) = w_1x_2(t)$. We can further deduce that necessarily $w_2 = w_3 = w_4 = w_6 = w_7 = 0$, and $w_5 = -w_1q$. In matrix form, this is equivalent to

$$\beta^\top YA(\kappa) = w_1(1, 0, 0, -q, 0, 0, 0, 0, 0, 0),$$

but this is not possible because in this case the vector on the right-hand side does not belong to the left image of $YA(\kappa)$.

### F Applications to the systems introduced in the main text.

Here we use the theory developed in this work to analyze the reaction systems introduced in the main text.

#### F.1 Toy example

Consider the mass-action system

$$A + B \xrightarrow{\kappa_1} 2B$$
$$B \xrightarrow{\kappa_2} A$$

Order the species alphabetically, and the complexes in the appearance order from left to right and from top to bottom, as $A + B$, $2B$, $B$, and $A$. In this case, we have

$$Y = \begin{pmatrix} 1 & 0 & 0 & 1 \\ 1 & 2 & 1 & 0 \end{pmatrix} \quad \text{and} \quad A(\kappa) = \begin{pmatrix} -\kappa_1 & 0 & 0 & 0 \\ \kappa_1 & 0 & 0 & 0 \\ 0 & 0 & -\kappa_2 & 0 \\ 0 & 0 & \kappa_2 & 0 \end{pmatrix} \tag{F.1}$$

The two non-terminal complexes differing for the entry relative to $A$ are the first and the third ones. Since the model has deficiency 1, it follows from Theorem 4.1 that $\hat{\Gamma}_{13}(\kappa)$ is non-empty. We can use Matlab to explicitly calculate it. First, in order to define $Y$ and $A(\kappa)$ in Matlab, we define the symbolic variables $\kappa_1$ and $\kappa_2$ and require they are positive real numbers via

```
k1=sym('k1','positive');
k2=sym('k2','positive');
```

We then define the matrices `Y` and `Ak` as in (F.1). We can calculate $\Psi_{13}(\kappa)$ via

```
e=eye(4);
simplify(null([Ak'*Y' e(:,1) e(:,3)]))
```

which returns the following matrix, whose columns are a basis for $\Psi_{13}(\kappa)$:

$$\begin{pmatrix} 1 & -\frac{1}{\kappa_2} \\ 1 & 0 \\ 0 & -\frac{\kappa_1}{\kappa_2} \\ 0 & 1 \end{pmatrix}.$$

It then follows from Proposition C.1 that

$$\Gamma_{13}(\kappa) = \left\{ \begin{pmatrix} \frac{1}{\kappa_2} \\ 0 \\ \frac{\kappa_1}{\kappa_2} \end{pmatrix} + a \begin{pmatrix} 1 \\ 1 \\ 0 \end{pmatrix} : a \in \mathbb{R} \right\},$$

which in turn implies that

$$\hat{\Gamma}_{13}(\kappa) = \left\{ \begin{pmatrix} \frac{1}{\kappa_2} \\ 0 \end{pmatrix} + a \begin{pmatrix} 1 \\ 1 \end{pmatrix} \ : \ a \in \mathbb{R} \right\} \tag{F.2}$$

and together with Theorem 4.2 that the ACR value of $A$ is $\kappa_1/\kappa_2$. Note that in this case

$$\mathcal{S} = \text{span}_{\mathbb{R}} \begin{pmatrix} 1 \\ -1 \end{pmatrix} \quad \text{and} \quad \mathcal{S}^{\perp} = \text{span}_{\mathbb{R}} \begin{pmatrix} 1 \\ 1 \end{pmatrix}. \tag{F.3}$$

Moreover, each connected component contains exactly one terminal component. Hence, Proposition C.2 applies, and as a matter of fact

$$\hat{\Gamma}_{13}(\kappa) = \begin{pmatrix} \frac{1}{\kappa_2} \\ 0 \end{pmatrix} + \mathcal{S}^{\perp}.$$

Finally, while it is clear from (F.2) that a vector with zero first component and a vector with zero second component are in $\hat{\Gamma}_{13}(\kappa)$, this could be derived from Corollary C.3 and from (F.3) without explicitly calculating $\hat{\Gamma}_{13}(\kappa)$. As a consequence, due to Theorem 6.1, disturbances could be applied to the production and degradation rates of $A$ (or $B$) while maintaining the absolute concentration robustness of the species $A$, its ACR value, and the linear CIF described in Theorem 4.2.

#### F.2 EnvZ-OmpR osmoregulatory system

Consider the mass-action system

$$\text{EnvZ-D} \underset{\kappa_2[\text{ADP}]}{\overset{\kappa_1}{\rightleftharpoons}} \text{EnvZ} \underset{\kappa_4}{\overset{\kappa_3[\text{ATP}]}{\rightleftharpoons}} \text{EnvZ-T} \xrightarrow{\kappa_5} \text{EnvZ-P}$$

$$\text{EnvZ-P} + \text{OmpR} \underset{\kappa_7}{\overset{\kappa_6}{\rightleftharpoons}} \text{EnvZ-OmpR-P} \xrightarrow{\kappa_8} \text{EnvZ} + \text{OmpR-P}$$

$$\text{EnvZ-D} + \text{OmpR-P} \underset{\kappa_{10}}{\overset{\kappa_9}{\rightleftharpoons}} \text{EnvZ-OmpR-D-P} \xrightarrow{\kappa_{11}} \text{EnvZ-D} + \text{OmpR}$$

Order both the species and the complexes according to their appearance order, from left to right and from top to bottom. Hence, the 8 species are ordered as EnvZ-D, EnvZ, EnvZ-T, EnvZ-P, OmpR, EnvZ-OmpR-P, OmpR-P, and EnvZ-OmpR-D-P. The 10 complexes are ordered as EnvZ-D, EnvZ, EnvZ-T, EnvZ-P, EnvZ-P+OmpR, EnvZ-OmpR-P, EnvZ+OmpR-P, EnvZ-D+OmpR-P, EnvZ-OmpR-D-P, and EnvZ-D+OmpR.

It can be checked that $\dim \mathcal{S} = 6$ and

$$\mathcal{S}^{\perp} = \text{span}_{\mathbb{R}} \left\{ \begin{pmatrix} 1 \\ 1 \\ 1 \\ 1 \\ 0 \\ 1 \\ 0 \\ 1 \end{pmatrix}, \begin{pmatrix} 0 \\ 0 \\ 0 \\ 0 \\ 1 \\ 1 \\ 1 \\ 1 \end{pmatrix} \right\}. \tag{F.4}$$

The above conservation laws correspond to the fact that the total mass of the chemical species containing some form of EnvZ is conserved, as well as the total mass of the chemical species containing some form of OmpR. We can calculate the deficiency as

$$\delta = m - \ell - \dim \mathcal{S} = 10 - 3 - 6 = 1.$$

The non-terminal complexes that only differ for the entry relative to OmpR-P are the first and the eighth ones. Therefore, we are interested in the study of $\hat{\Gamma}_{18}(\kappa)$, which by Theorem 4.1 is not empty. We have

$$Y = \begin{pmatrix} 1 & 0 & 0 & 0 & 0 & 0 & 0 & 1 & 0 & 1 \\ 0 & 1 & 0 & 0 & 0 & 0 & 1 & 0 & 0 & 0 \\ 0 & 0 & 1 & 0 & 0 & 0 & 0 & 0 & 0 & 0 \\ 0 & 0 & 0 & 1 & 1 & 0 & 0 & 0 & 0 & 0 \\ 0 & 0 & 0 & 0 & 1 & 0 & 0 & 0 & 0 & 1 \\ 0 & 0 & 0 & 0 & 0 & 1 & 0 & 0 & 0 & 0 \\ 0 & 0 & 0 & 0 & 0 & 0 & 1 & 1 & 0 & 0 \\ 0 & 0 & 0 & 0 & 0 & 0 & 0 & 0 & 1 & 0 \end{pmatrix}$$

and

$$A(\kappa)=\begin{pmatrix}-\kappa_1 & \kappa_2[\mathrm{ADP}] & 0 & 0 & 0 & 0 & 0 & 0 & 0 & 0\\ \kappa_1 & -\kappa_2[\mathrm{ADP}]-\kappa_3[\mathrm{ATP}] & \kappa_4 & 0 & 0 & 0 & 0 & 0 & 0 & 0\\ 0 & \kappa_3[\mathrm{ATP}] & -\kappa_4-\kappa_5 & 0 & 0 & 0 & 0 & 0 & 0 & 0\\ 0 & 0 & \kappa_5 & 0 & 0 & 0 & 0 & 0 & 0 & 0\\ 0 & 0 & 0 & 0 & -\kappa_6 & \kappa_7 & 0 & 0 & 0 & 0\\ 0 & 0 & 0 & 0 & \kappa_6 & -\kappa_7-\kappa_8 & 0 & 0 & 0 & 0\\ 0 & 0 & 0 & 0 & 0 & \kappa_8 & 0 & 0 & 0 & 0\\ 0 & 0 & 0 & 0 & 0 & 0 & 0 & -\kappa_9 & \kappa_{10} & 0\\ 0 & 0 & 0 & 0 & 0 & 0 & 0 & \kappa_9 & -\kappa_{10}-\kappa_{11} & 0\\ 0 & 0 & 0 & 0 & 0 & 0 & 0 & 0 & \kappa_{11} & 0\end{pmatrix}.$$

In order to define the corresponding symbolic matrices `Y` and `Ak` in Matlab, we can first define the symbolic positive real variables

```
k=sym('k', [1,11], 'positive');
k(2)=k(2)*sym('ADP', 'positive');
k(3)=k(3)*sym('ATP', 'positive');
```

Then we calculate $\Psi_{18}(\kappa)$, as defined in (C.3), via

```
e=eye(10);
simplify(null([Ak'*Y' e(:,8) e(:,1)]))
```

The output of the last command is the following matrix, whose columns are a basis of $\Psi_{18}(\kappa)$.

$$\begin{pmatrix}-1 & 1 & -\frac{[\mathrm{ADP}]\,\kappa_2\,\kappa_{11}\,(\kappa_4+\kappa_5)}{[\mathrm{ATP}]\,\kappa_1\,\kappa_3\,\kappa_5\,(\kappa_{10}+\kappa_{11})}\\ -1 & 1 & -\frac{[\mathrm{ADP}]\,\kappa_2\,\kappa_4\,\kappa_{11}+[\mathrm{ADP}]\,\kappa_2\,\kappa_5\,\kappa_{11}+[\mathrm{ATP}]\,\kappa_3\,\kappa_5\,\kappa_{10}+[\mathrm{ATP}]\,\kappa_3\,\kappa_5\,\kappa_{11}}{[\mathrm{ATP}]\,\kappa_1\,\kappa_3\,\kappa_5\,(\kappa_{10}+\kappa_{11})}\\ -1 & 1 & -\frac{[\mathrm{ADP}]\,\kappa_2\,\kappa_4\,\kappa_{11}+[\mathrm{ADP}]\,\kappa_2\,\kappa_5\,\kappa_{10}+2\,[\mathrm{ADP}]\,\kappa_2\,\kappa_5\,\kappa_{11}+[\mathrm{ATP}]\,\kappa_3\,\kappa_5\,\kappa_{10}+[\mathrm{ATP}]\,\kappa_3\,\kappa_5\,\kappa_{11}}{[\mathrm{ATP}]\,\kappa_1\,\kappa_3\,\kappa_5\,(\kappa_{10}+\kappa_{11})}\\ -1 & 1 & -\frac{[\mathrm{ADP}]\,\kappa_2\,\kappa_4\,\kappa_{10}+2\,[\mathrm{ADP}]\,\kappa_2\,\kappa_4\,\kappa_{11}+[\mathrm{ADP}]\,\kappa_2\,\kappa_5\,\kappa_{10}+2\,[\mathrm{ADP}]\,\kappa_2\,\kappa_5\,\kappa_{11}+[\mathrm{ATP}]\,\kappa_3\,\kappa_5\,\kappa_{10}+[\mathrm{ATP}]\,\kappa_3\,\kappa_5\,\kappa_{11}}{[\mathrm{ATP}]\,\kappa_1\,\kappa_3\,\kappa_5\,(\kappa_{10}+\kappa_{11})}\\ 1 & 0 & \frac{[\mathrm{ADP}]\,\kappa_2\,(\kappa_4+\kappa_5)}{[\mathrm{ATP}]\,\kappa_1\,\kappa_3\,\kappa_5}\\ 0 & 1 & -\frac{[\mathrm{ADP}]\,\kappa_2\,\kappa_4\,\kappa_{11}+[\mathrm{ADP}]\,\kappa_2\,\kappa_5\,\kappa_{11}+[\mathrm{ATP}]\,\kappa_3\,\kappa_5\,\kappa_{10}+[\mathrm{ATP}]\,\kappa_3\,\kappa_5\,\kappa_{11}}{[\mathrm{ATP}]\,\kappa_1\,\kappa_3\,\kappa_5\,(\kappa_{10}+\kappa_{11})}\\ 1 & 0 & 0\\ 0 & 1 & 0\\ 0 & 0 & -\frac{[\mathrm{ADP}]\,\kappa_2\,\kappa_9\,\kappa_{11}\,(\kappa_4+\kappa_5)}{[\mathrm{ATP}]\,\kappa_1\,\kappa_3\,\kappa_5\,(\kappa_{10}+\kappa_{11})}\\ 0 & 0 & 1\end{pmatrix}$$

For convenience, denote the last vector by $\zeta(\kappa)$. It follows from Proposition C.1 that

$$\Gamma_{18}(\kappa)=\left\{-\pi_9(\zeta(\kappa))+a_1\begin{pmatrix}-1\\-1\\-1\\-1\\1\\0\\1\\0\\0\end{pmatrix}+a_2\begin{pmatrix}1\\1\\1\\1\\0\\1\\0\\1\\0\end{pmatrix}\ :\ a_1,a_2\in\mathbb{R}\right\}.$$

It follows that

$$\hat{\Gamma}_{18}(\kappa) = \left\{ -\pi_8(\zeta(\kappa)) + a_1 \begin{pmatrix} -1 \\ -1 \\ -1 \\ -1 \\ 1 \\ 0 \\ 1 \\ 0 \end{pmatrix} + a_2 \begin{pmatrix} 1 \\ 1 \\ 1 \\ 1 \\ 0 \\ 1 \\ 0 \\ 1 \end{pmatrix} \; : \; a_1, a_2 \in \mathbb{R} \right\}, \tag{F.5}$$

which corresponds to (11) in the main text. Moreover, it follows from Theorem 4.2 that the ACR value of the species OmpR-P is

$$-\zeta_9(\kappa) = \frac{[\mathrm{ADP}]\,\kappa_2\,\kappa_9\,\kappa_{11}\,\left(\kappa_4 + \kappa_5\right)}{[\mathrm{ATP}]\,\kappa_1\,\kappa_3\,\kappa_5\,\left(\kappa_{10} + \kappa_{11}\right)}$$

as reported in (10) in the main text.

Further things can be noted about this model. First, it follows from (F.4) and (F.5) that

$$\hat{\Gamma}_{18}(\kappa) = -\pi_8(\zeta(\kappa)) + \mathcal{S}^{\perp},$$

which is in accordance with Proposition C.2 because all connected components contain exactly one terminal component. Secondly, due to Corollary C.3, we can deduce without explicitly calculating $\Gamma_{18}(\kappa)$ that there is a vector $\hat{\gamma}$ in $\hat{\Gamma}_{18}(\kappa)$ whose entries relative to species EnvZ and OmpR-P are zero (which are the second and the seventh complexes, respectively). Specifically, if we consider the basis of $\mathcal{S}^{\perp}$ given in (F.4) and we let $X_{l_1} = X_2 =$EnvZ and $X_{l_2} = X_7 =$OmpR-P, we have

$$V = \begin{pmatrix} 1 & 0 \\ 0 & 1 \end{pmatrix},$$

which has rank 2. The existence of such vector $\hat{\gamma}$ is used in Section 6.3 and it implies by Theorem 6.1 that the system is robust to persistent disturbances affecting the production and degradation rates of both EnvZ and OmpR-P.

### G An ACR signaling system covered by our theory and not by [40]

Consider the double-phosphorylation mass-action system in Figure 6. We will show that the theory developed in [40] stays silent on whether it is ACR. However, our theory covers this case and implies the double-phosphorylated form of the transcriptional regulatory protein is ACR, for any choice of kinetic parameters $\kappa$ such that a positive steady state exists. Note that this form of response robustness may seem a bit surprising for a multisite phosphorylation mechanism, since these are often known for their multi-stability properties, notably shown in the case of the MAPK pathway [7, 26, 27, 44].

Order the species and the complexes according to their appearance order, from left to right and from top to bottom. The 11 species are then ordered as $A$, $A^{\star}$, $A$-P, $B$, $A$-$B$-P, $B$-P, $A$-$B$-PP, $B$-PP, $A^{\star}$-$B$-P, $A^{\star}$-$B$-PP, and $B$-PP-$A$. The 13 complexes are ordered as $A$, $A^{\star}$, $A$-P, $A$-P + $B$, $A$-$B$-P, $A$ + $B$-P, $A$-P + $B$-P, $A$-$B$-PP, $A$ + $B$-PP, $A^{\star}$-$B$-P, $A^{\star}$-$B$-PP, $A$-P + $B$-PP, and $B$-PP-$A$. It can be checked that $\dim \mathcal{S} = 9$ and

$$\mathcal{S}^{\perp} = \mathrm{span}_{\mathbb{R}} \left\{ \begin{pmatrix} 1 \\ 1 \\ 1 \\ 0 \\ 1 \\ 0 \\ 1 \\ 0 \\ 1 \\ 1 \\ 1 \end{pmatrix}, \begin{pmatrix} 0 \\ 0 \\ 0 \\ 1 \\ 1 \\ 1 \\ 1 \\ 1 \\ 1 \\ 1 \\ 1 \end{pmatrix} \right\}. \tag{G.1}$$

Similarly as for the EnvZ-OmpR osmoregulatory system, the above conservation laws correspond express that the total mass of the chemical species containing some form of the protein $A$ is conserved, as well as the total mass of the chemical species containing some form of the protein $B$. The reaction graph is given in Figure 7. In particular, $\ell = 2$ and the deficiency is

$$\delta = m - \ell - \dim \mathcal{S} = 13 - 2 - 9 = 2.$$

So, this model is not included in the class of models studied in [40], and Theorems 3.1 and B.3 do not apply.

Activation and phosphorylation of the sensor-transmitter protein:

$$A \underset{\kappa_2}{\overset{\kappa_1[\mathrm{I}]}{\rightleftharpoons}} A^{\star} \xrightarrow{\kappa_3} A\text{-}\mathbf{P}$$

Phosphorylation of the sensory response protein:

$$A\text{-}\mathbf{P} + B \underset{\kappa_5}{\overset{\kappa_4}{\rightleftharpoons}} A\text{-}B\text{-}\mathbf{P} \xrightarrow{\kappa_6} A + B\text{-}\mathbf{P}$$

$$A\text{-}\mathbf{P} + B\text{-}\mathbf{P} \underset{\kappa_8}{\overset{\kappa_7}{\rightleftharpoons}} A\text{-}B\text{-}\mathbf{PP} \xrightarrow{\kappa_9} A + B\text{-}\mathbf{PP}$$

Activation and phosphorylation of the protein aggregate:

$$A\text{-}B\text{-}\mathbf{P} \underset{\kappa_{11}}{\overset{\kappa_{10}[\mathrm{I}]}{\rightleftharpoons}} A^{\star}\text{-}B\text{-}\mathbf{P}$$

$$A\text{-}B\text{-}\mathbf{PP} \underset{\kappa_{13}}{\overset{\kappa_{12}[\mathrm{I}]}{\rightleftharpoons}} A^{\star}\text{-}B\text{-}\mathbf{PP}$$

$$A^{\star}\text{-}B\text{-}\mathbf{P} \xrightarrow{\kappa_{14}} A^{\star}\text{-}B\text{-}\mathbf{PP} \xrightarrow{\kappa_{15}} A\text{-}\mathbf{P} + B\text{-}\mathbf{PP}$$

Dephosphorylation of the sensory response protein:

$$A + B\text{-}\mathbf{PP} \underset{\kappa_{17}}{\overset{\kappa_{16}}{\rightleftharpoons}} B\text{-}\mathbf{PP}\text{-}A \xrightarrow{\kappa_{18}} A\text{-}\mathbf{P} + B$$

Figure 6: A model of signal transduction. Here, the sensory response protein $B$ is active if two phosphoryl groups are attached. In the first line, the protein $A$ is phosphorylated in response to a stimulus [I]. In the second and third line, the phosphoryl groups are transferred to the sensory response protein $B$. In the forth and fifth line, the protein $A$ responds by the stimulus [I] while it is bound to $B$. The resulting protein aggregate is able to gain phosphoryl groups until the three phosphoryl sites are occupied (as described in the sixth line). Finally, in the last two lines, the inactive form of the sensor-transmitter protein $A$ acts as a phosphatase on $B$.

The first and the ninth complexes are non-terminal, and they only differ for the entry relative B-PP. Hence, in order to apply Theorems 4.2 and 6.1, we are interested in studying $\hat{\Gamma}_{19}(\kappa)$. Since the deficiency of the model is 2, Theorem 4.1 does not apply, so to understand whether $\hat{\Gamma}_{19}(\kappa)$ is non-empty we need to explicitly calculate it. To this aim, we define in Matlab the following positive symbolic variables, which correspond to the rate constants of the model:

```
k=sym('k', [1,18], 'positive');
k(1)=k(1)*sym('I', 'positive');
k(10)=k(10)*sym('I', 'positive');
k(12)=k(12)*sym('I', 'positive');
```

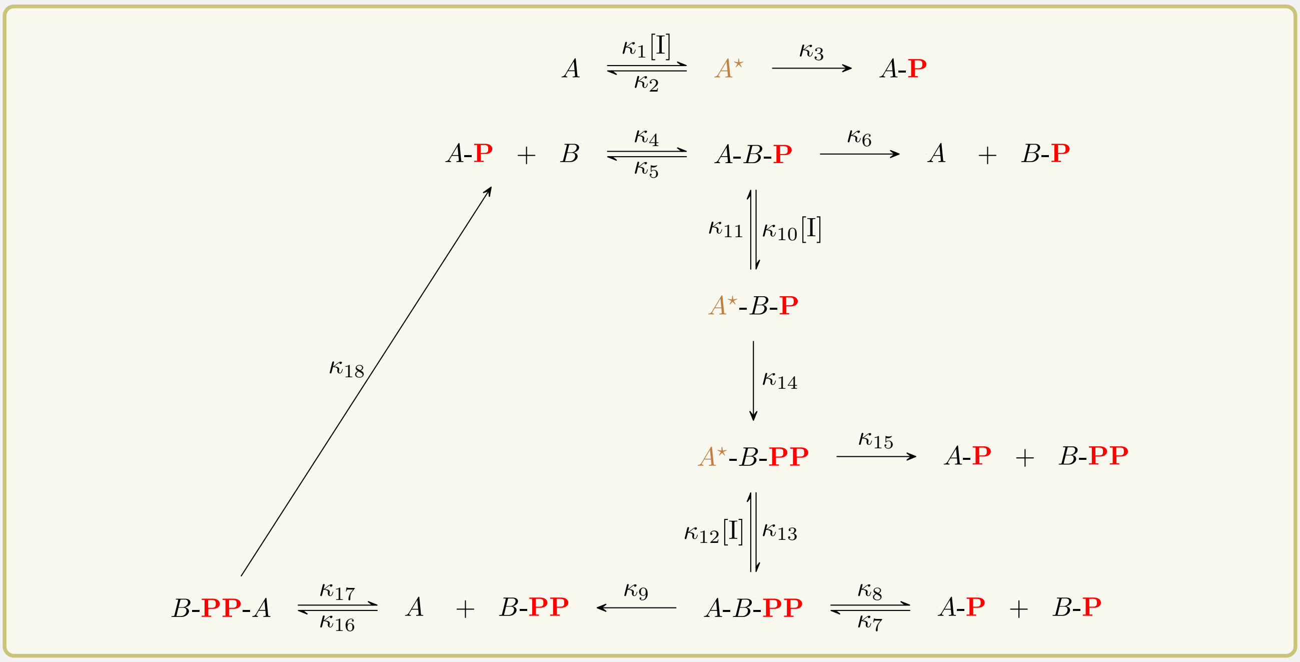


Figure 7: Reaction graph for the signaling transduction system proposed in Figure 6.

We then define the symbolic matrices `Y` and `Ak` corresponding to

$$Y = \begin{pmatrix} 1 & 0 & 0 & 0 & 0 & 1 & 0 & 0 & 1 & 0 & 0 & 0 & 0 \\ 0 & 1 & 0 & 0 & 0 & 0 & 0 & 0 & 0 & 0 & 0 & 0 & 0 \\ 0 & 0 & 1 & 1 & 0 & 0 & 1 & 0 & 0 & 0 & 0 & 1 & 0 \\ 0 & 0 & 0 & 1 & 0 & 0 & 0 & 0 & 0 & 0 & 0 & 0 & 0 \\ 0 & 0 & 0 & 0 & 1 & 0 & 0 & 0 & 0 & 0 & 0 & 0 & 0 \\ 0 & 0 & 0 & 0 & 0 & 1 & 1 & 0 & 0 & 0 & 0 & 0 & 0 \\ 0 & 0 & 0 & 0 & 0 & 0 & 0 & 1 & 0 & 0 & 0 & 0 & 0 \\ 0 & 0 & 0 & 0 & 0 & 0 & 0 & 0 & 1 & 0 & 0 & 1 & 0 \\ 0 & 0 & 0 & 0 & 0 & 0 & 0 & 0 & 0 & 1 & 0 & 0 & 0 \\ 0 & 0 & 0 & 0 & 0 & 0 & 0 & 0 & 0 & 0 & 1 & 0 & 0 \\ 0 & 0 & 0 & 0 & 0 & 0 & 0 & 0 & 0 & 0 & 0 & 0 & 1 \end{pmatrix}$$

and to $A(\kappa)$ as described in Figure 8, respectively.

In order to calculate a basis for $\Psi_{19}(\kappa)$ as defined in (C.3), we type

```
e=eye(13);
simplify(null([Ak'*Y' e(:,1) e(:,9)]))
```

The output of the last command is shown in Figure 9. While the output is rather complicated, we did not need to put much effort in its calculation, which was completed in Matlab in a matter of seconds. For convenience we denote by $\zeta(\kappa)$ the last column of the output matrix.

It follows from Proposition C.1 that $\Gamma_{19}(\kappa)$ is non-empty, and is given by

$$\Gamma_{19}(\kappa) = \left\{ -\pi_{12}(\zeta(\kappa)) + a_1 \begin{pmatrix} -1 \\ -1 \\ -1 \\ 1 \\ 0 \\ 1 \\ 0 \\ 1 \\ 0 \\ 0 \\ 0 \\ 0 \end{pmatrix} + a_2 \begin{pmatrix} 1 \\ 1 \\ 1 \\ 0 \\ 1 \\ 0 \\ 1 \\ 0 \\ 1 \\ 1 \\ 1 \\ 0 \end{pmatrix} : a_1, a_2 \in \mathbb{R} \right\}. \tag{G.2}$$

It follows from (G.2) and (G.1) that

$$\hat{\Gamma}_{19}(\kappa) = \left\{ -\pi_{11}(\zeta(\kappa)) + a_1 \begin{pmatrix} -1 \\ -1 \\ -1 \\ 1 \\ 0 \\ 1 \\ 0 \\ 1 \\ 0 \\ 0 \\ 0 \end{pmatrix} + a_2 \begin{pmatrix} 1 \\ 1 \\ 1 \\ 0 \\ 1 \\ 0 \\ 1 \\ 0 \\ 1 \\ 1 \\ 1 \end{pmatrix} : a_1, a_2 \in \mathbb{R} \right\} = -\pi_{11}(\zeta(\kappa)) + \mathcal{S}^{\perp},$$

which is in accordance with Proposition C.2 because every connected component of the reaction graph in Figure 7 contains exactly one terminal component. Moreover, from (G.2) and Theorem 4.2 it follows that the ACR value of the ACR species $B$-PP is

$$-\zeta_{12}(\kappa) = \frac{\kappa_1\kappa_3[\mathrm{I}](\kappa_{17}+\kappa_{18})(\kappa_6\kappa_{11}+\kappa_6\kappa_{14}+\kappa_{10}\kappa_{14}[\mathrm{I}])(\kappa_9\kappa_{13}+\kappa_9\kappa_{15}+\kappa_{12}\kappa_{15}[\mathrm{I}])}{\kappa_{16}\kappa_{18}(\kappa_2+\kappa_3)g(\kappa,[\mathrm{I}])},$$

where

$$g(\kappa,[\mathrm{I}]) = -\kappa_{10}\kappa_{12}\kappa_{14}\kappa_{15}[\mathrm{I}]^2 - \kappa_9\kappa_{10}\kappa_{14}\kappa_{15}[\mathrm{I}] + \kappa_6\kappa_9\kappa_{11}\kappa_{13} + \kappa_6\kappa_9\kappa_{11}\kappa_{15} + \kappa_6\kappa_9\kappa_{13}\kappa_{14} + \kappa_6\kappa_9\kappa_{14}\kappa_{15}.$$

This means that if a positive steady state $c$ exists, necessarily it entry relative to $B$-PP (which is the eighth species) satisfies $c_8 = -\zeta_{12}(\kappa)$. Of course, this cannot occur if $-\zeta_{12}(\kappa)$ is non-positive, i.e. if $g(\kappa,[\mathrm{I}])$ is non-positive, in which case no positive steady state exists. Note that we did not need to work directly with the differential equation to derive this information. We will further show that in fact a positive steady state exists if and only if $g(\kappa,[\mathrm{I}])$ is positive. To this aim, note that $c$ is a steady state if and only if $\Lambda(c) \in \ker YA(\kappa)$. Then, we calculate a basis for $\ker YA(\kappa)$ by typing

```
simplify(null(Y*Ak))
```

which returns a matrix of the form

$$\begin{pmatrix} 0 & 0 & 0 & b_1(\kappa,[\mathrm{I}])g(\kappa,[\mathrm{I}]) \\ 0 & 0 & 0 & b_2(\kappa,[\mathrm{I}])g(\kappa,[\mathrm{I}]) \\ 1 & 0 & 0 & 0 \\ 0 & 0 & 0 & b_4(\kappa,[\mathrm{I}]) \\ 0 & 0 & 0 & b_5(\kappa,[\mathrm{I}]) \\ 0 & 1 & 0 & 0 \\ 0 & 0 & 0 & b_7(\kappa,[\mathrm{I}]) \\ 0 & 0 & 0 & b_8(\kappa,[\mathrm{I}]) \\ 0 & 0 & 0 & b_9(\kappa,[\mathrm{I}]) \\ 0 & 0 & 0 & b_{10}(\kappa,[\mathrm{I}]) \\ 0 & 0 & 0 & b_{11}(\kappa,[\mathrm{I}]) \\ 0 & 0 & 1 & 0 \\ 0 & 0 & 0 & 1 \end{pmatrix}$$

for some functions $b_i(\kappa, [\mathrm{I}])$ which map positive arguments to positive real numbers. Hence, a positive steady state exists if and only if there exists a positive vector $c$ such that

$$
\begin{aligned}
c_1 &= a_4 b_1(\kappa,[\mathrm{I}])g(\kappa,[\mathrm{I}]) & c_1c_6 &= a_2 & c_9 &= a_4 b_{10}(\kappa,[\mathrm{I}])\\
c_2 &= a_4 b_2(\kappa,[\mathrm{I}])g(\kappa,[\mathrm{I}]) & c_3c_6 &= a_4 b_7(\kappa,[\mathrm{I}]) & c_{10} &= a_4 b_{11}(\kappa,[\mathrm{I}])\\
c_3 &= a_1 & c_7 &= a_4 b_8(\kappa,[\mathrm{I}]) & c_3c_8 &= a_3\\
c_3c_4 &= a_4 b_4(\kappa,[\mathrm{I}]) & c_1c_8 &= a_4 b_9(\kappa,[\mathrm{I}]) & c_{11} &= a_4\\
c_5 &= a_4 b_5(\kappa,[\mathrm{I}])
\end{aligned}
$$

for some $a_1, a_2, a_3, a_4 \in \mathbb{R}_{>0}$. If $g(\kappa, [\mathrm{I}])$ is positive, it is easy to see that such a positive vector $c$ exists. Specifically, the positive steady states are parameterized by

$$
\begin{aligned}
c_1 &= a_4 b_1(\kappa,[\mathrm{I}])g(\kappa,[\mathrm{I}]) & c_5 &= a_4 b_5(\kappa,[\mathrm{I}]) & c_9 &= a_4 b_{10}(\kappa,[\mathrm{I}])\\
c_2 &= a_4 b_2(\kappa,[\mathrm{I}])g(\kappa,[\mathrm{I}]) & c_6 &= \frac{a_4}{a_1} b_7(\kappa,[\mathrm{I}]) & c_{10} &= a_4 b_{11}(\kappa,[\mathrm{I}])\\
c_3 &= a_1 & c_7 &= a_4 b_8(\kappa,[\mathrm{I}]) & c_{11} &= a_4\\
c_4 &= \frac{a_4}{a_1} b_4(\kappa,[\mathrm{I}]) & c_8 &= \frac{b_9(\kappa,[\mathrm{I}])}{b_1(\kappa,[\mathrm{I}])g(\kappa,[\mathrm{I}])}
\end{aligned}
$$

where $a_1$ and $a_4$ vary in $\mathbb{R}_{>0}$. Note that the entry $c_8$ is relative to the ACR species $B$-PP and is fixed.

$$A(\kappa) = \begin{pmatrix}
-\kappa_1[\mathrm{I}] & \kappa_2 & 0 & 0 & 0 & 0 & 0 & 0 & 0 & 0 & 0 & 0 & 0 \\
\kappa_1[\mathrm{I}] & -\kappa_2-\kappa_3 & 0 & 0 & 0 & 0 & 0 & 0 & 0 & 0 & 0 & 0 & 0 \\
0 & \kappa_3 & 0 & 0 & 0 & 0 & 0 & 0 & 0 & 0 & 0 & 0 & 0 \\
0 & 0 & 0 & -\kappa_4 & \kappa_5 & 0 & 0 & 0 & 0 & 0 & 0 & 0 & \kappa_{18} \\
0 & 0 & 0 & \kappa_4 & -\kappa_5-\kappa_6-\kappa_{10}[\mathrm{I}] & 0 & 0 & 0 & 0 & \kappa_{11} & 0 & 0 & 0 \\
0 & 0 & 0 & 0 & \kappa_6 & 0 & 0 & 0 & 0 & 0 & 0 & 0 & 0 \\
0 & 0 & 0 & 0 & 0 & 0 & -\kappa_7 & \kappa_8 & 0 & 0 & 0 & 0 & 0 \\
0 & 0 & 0 & 0 & 0 & 0 & \kappa_7 & -\kappa_8-\kappa_9-\kappa_{12}[\mathrm{I}] & 0 & 0 & \kappa_{13} & 0 & 0 \\
0 & 0 & 0 & 0 & 0 & 0 & 0 & \kappa_9 & -\kappa_{16} & 0 & 0 & 0 & \kappa_{17} \\
0 & 0 & 0 & 0 & \kappa_{10}[\mathrm{I}] & 0 & 0 & 0 & 0 & -\kappa_{11}-\kappa_{14} & 0 & 0 & 0 \\
0 & 0 & 0 & 0 & 0 & 0 & 0 & \kappa_{12}[\mathrm{I}] & 0 & \kappa_{14} & -\kappa_{13}-\kappa_{15} & 0 & 0 \\
0 & 0 & 0 & 0 & 0 & 0 & 0 & 0 & 0 & 0 & \kappa_{15} & 0 & 0 \\
0 & 0 & 0 & 0 & 0 & 0 & 0 & 0 & \kappa_{16} & 0 & 0 & 0 & -\kappa_{17}-\kappa_{18}
\end{pmatrix}$$

Figure 8: $A(k)$ for the mass-action system in Figure 6.

$$
\begin{pmatrix}
-1 & 1 & \frac{1}{\kappa_{16}} \\
-1 & 1 & \frac{p_1(\kappa,[\mathrm{I}])}{\kappa_{16}\kappa_{18}(\kappa_2+\kappa_3)\left(-\kappa_{10}\kappa_{12}\kappa_{14}\kappa_{15}[\mathrm{I}]^2-\kappa_9\kappa_{10}\kappa_{14}\kappa_{15}[\mathrm{I}]+\kappa_6\kappa_9\kappa_{11}\kappa_{13}+\kappa_6\kappa_9\kappa_{11}\kappa_{15}+\kappa_6\kappa_9\kappa_{13}\kappa_{14}+\kappa_6\kappa_9\kappa_{14}\kappa_{15}\right)} \\
-1 & 1 & \frac{p_2(\kappa,[\mathrm{I}])}{\kappa_{16}\kappa_{18}\left(-\kappa_{10}\kappa_{12}\kappa_{14}\kappa_{15}[\mathrm{I}]^2-\kappa_9\kappa_{10}\kappa_{14}\kappa_{15}[\mathrm{I}]+\kappa_6\kappa_9\kappa_{11}\kappa_{13}+\kappa_6\kappa_9\kappa_{11}\kappa_{15}+\kappa_6\kappa_9\kappa_{13}\kappa_{14}+\kappa_6\kappa_9\kappa_{14}\kappa_{15}\right)} \\
1 & 0 & -\frac{(\kappa_{17}+\kappa_{18})(2\kappa_6\kappa_9\kappa_{11}\kappa_{13}+2\kappa_6\kappa_9\kappa_{11}\kappa_{15}+2\kappa_6\kappa_9\kappa_{13}\kappa_{14}+2\kappa_6\kappa_9\kappa_{14}\kappa_{15}+[\mathrm{I}]\kappa_6\kappa_{11}\kappa_{12}\kappa_{15}+[\mathrm{I}]\kappa_9\kappa_{10}\kappa_{13}\kappa_{14}+[\mathrm{I}]\kappa_6\kappa_{12}\kappa_{14}\kappa_{15})}{\kappa_{16}\kappa_{18}\left(-\kappa_{10}\kappa_{12}\kappa_{14}\kappa_{15}[\mathrm{I}]^2-\kappa_9\kappa_{10}\kappa_{14}\kappa_{15}[\mathrm{I}]+\kappa_6\kappa_9\kappa_{11}\kappa_{13}+\kappa_6\kappa_9\kappa_{11}\kappa_{15}+\kappa_6\kappa_9\kappa_{13}\kappa_{14}+\kappa_6\kappa_9\kappa_{14}\kappa_{15}\right)} \\
0 & 1 & -\frac{\kappa_{17}}{\kappa_{16}\kappa_{18}} \\
1 & 0 & -\frac{\kappa_9(\kappa_{13}+\kappa_{15})(\kappa_{17}+\kappa_{18})(\kappa_6\kappa_{11}+\kappa_6\kappa_{14}+[\mathrm{I}]\kappa_{10}\kappa_{14})}{\kappa_{16}\kappa_{18}\left(-\kappa_{10}\kappa_{12}\kappa_{14}\kappa_{15}[\mathrm{I}]^2-\kappa_9\kappa_{10}\kappa_{14}\kappa_{15}[\mathrm{I}]+\kappa_6\kappa_9\kappa_{11}\kappa_{13}+\kappa_6\kappa_9\kappa_{11}\kappa_{15}+\kappa_6\kappa_9\kappa_{13}\kappa_{14}+\kappa_6\kappa_9\kappa_{14}\kappa_{15}\right)} \\
0 & 1 & \frac{p_3(\kappa,[\mathrm{I}])}{\kappa_{16}\kappa_{18}\left(-\kappa_{10}\kappa_{12}\kappa_{14}\kappa_{15}[\mathrm{I}]^2-\kappa_9\kappa_{10}\kappa_{14}\kappa_{15}[\mathrm{I}]+\kappa_6\kappa_9\kappa_{11}\kappa_{13}+\kappa_6\kappa_9\kappa_{11}\kappa_{15}+\kappa_6\kappa_9\kappa_{13}\kappa_{14}+\kappa_6\kappa_9\kappa_{14}\kappa_{15}\right)} \\
1 & 0 & 0 \\
0 & 1 & \frac{\kappa_6\kappa_9\kappa_{13}\kappa_{14}\kappa_{18}-\kappa_6\kappa_9\kappa_{11}\kappa_{15}\kappa_{17}-\kappa_6\kappa_9\kappa_{11}\kappa_{13}\kappa_{17}+\kappa_6\kappa_9\kappa_{14}\kappa_{15}\kappa_{17}+2\kappa_6\kappa_9\kappa_{14}\kappa_{15}\kappa_{18}+[\mathrm{I}]^2\kappa_{10}\kappa_{12}\kappa_{14}\kappa_{15}\kappa_{17}+[\mathrm{I}]\kappa_6\kappa_{12}\kappa_{14}\kappa_{15}\kappa_{17}+[\mathrm{I}]\kappa_6\kappa_{12}\kappa_{14}\kappa_{15}\kappa_{18}+[\mathrm{I}]\kappa_9\kappa_{10}\kappa_{14}\kappa_{15}\kappa_{17}}{\kappa_{16}\kappa_{18}\left(-\kappa_{10}\kappa_{12}\kappa_{14}\kappa_{15}[\mathrm{I}]^2-\kappa_9\kappa_{10}\kappa_{14}\kappa_{15}[\mathrm{I}]+\kappa_6\kappa_9\kappa_{11}\kappa_{13}+\kappa_6\kappa_9\kappa_{11}\kappa_{15}+\kappa_6\kappa_9\kappa_{13}\kappa_{14}+\kappa_6\kappa_9\kappa_{14}\kappa_{15}\right)} \\
0 & 1 & \frac{p_4(\kappa,[\mathrm{I}])}{\kappa_{16}\kappa_{18}\left(-\kappa_{10}\kappa_{12}\kappa_{14}\kappa_{15}[\mathrm{I}]^2-\kappa_9\kappa_{10}\kappa_{14}\kappa_{15}[\mathrm{I}]+\kappa_6\kappa_9\kappa_{11}\kappa_{13}+\kappa_6\kappa_9\kappa_{11}\kappa_{15}+\kappa_6\kappa_9\kappa_{13}\kappa_{14}+\kappa_6\kappa_9\kappa_{14}\kappa_{15}\right)} \\
0 & 1 & 0 \\
0 & 0 & -\frac{[\mathrm{I}]\kappa_1\kappa_3(\kappa_{17}+\kappa_{18})(\kappa_6\kappa_{11}+\kappa_6\kappa_{14}+[\mathrm{I}]\kappa_{10}\kappa_{14})(\kappa_9\kappa_{13}+\kappa_9\kappa_{15}+[\mathrm{I}]\kappa_{12}\kappa_{15})}{\kappa_{16}\kappa_{18}(\kappa_2+\kappa_3)\left(-\kappa_{10}\kappa_{12}\kappa_{14}\kappa_{15}[\mathrm{I}]^2-\kappa_9\kappa_{10}\kappa_{14}\kappa_{15}[\mathrm{I}]+\kappa_6\kappa_9\kappa_{11}\kappa_{13}+\kappa_6\kappa_9\kappa_{11}\kappa_{15}+\kappa_6\kappa_9\kappa_{13}\kappa_{14}+\kappa_6\kappa_9\kappa_{14}\kappa_{15}\right)} \\
0 & 0 & 1
\end{pmatrix}
$$

where

$$
\begin{aligned}
p_1(\kappa,[\mathrm{I}]) = {} & \kappa_2\kappa_6\kappa_9\kappa_{11}\kappa_{13}\kappa_{18}+\kappa_3\kappa_6\kappa_9\kappa_{11}\kappa_{13}\kappa_{17}+2\kappa_3\kappa_6\kappa_9\kappa_{11}\kappa_{13}\kappa_{18}+\kappa_2\kappa_6\kappa_9\kappa_{11}\kappa_{15}\kappa_{18}+\kappa_3\kappa_6\kappa_9\kappa_{11}\kappa_{15}\kappa_{17}+\kappa_2\kappa_6\kappa_9\kappa_{13}\kappa_{14}\kappa_{18}+2\kappa_3\kappa_6\kappa_9\kappa_{11}\kappa_{15}\kappa_{18} \\
& +\kappa_3\kappa_6\kappa_9\kappa_{13}\kappa_{14}\kappa_{17}+2\kappa_3\kappa_6\kappa_9\kappa_{13}\kappa_{14}\kappa_{18}+\kappa_2\kappa_6\kappa_9\kappa_{14}\kappa_{15}\kappa_{18}+\kappa_3\kappa_6\kappa_9\kappa_{14}\kappa_{15}\kappa_{17}+2\kappa_3\kappa_6\kappa_9\kappa_{14}\kappa_{15}\kappa_{18}+[\mathrm{I}]\kappa_3\kappa_6\kappa_{11}\kappa_{12}\kappa_{15}\kappa_{17} \\
& +[\mathrm{I}]\kappa_3\kappa_6\kappa_{11}\kappa_{12}\kappa_{15}\kappa_{18}+[\mathrm{I}]\kappa_3\kappa_9\kappa_{10}\kappa_{13}\kappa_{14}\kappa_{17}+[\mathrm{I}]\kappa_3\kappa_6\kappa_{12}\kappa_{14}\kappa_{15}\kappa_{17}+[\mathrm{I}]\kappa_3\kappa_9\kappa_{10}\kappa_{13}\kappa_{14}\kappa_{18}-[\mathrm{I}]\kappa_2\kappa_9\kappa_{10}\kappa_{14}\kappa_{15}\kappa_{18}+[\mathrm{I}]\kappa_3\kappa_6\kappa_{12}\kappa_{14}\kappa_{15}\kappa_{18} \\
& +[\mathrm{I}]\kappa_3\kappa_9\kappa_{10}\kappa_{14}\kappa_{15}\kappa_{17}-[\mathrm{I}]^2\kappa_2\kappa_{10}\kappa_{12}\kappa_{14}\kappa_{15}\kappa_{18}+[\mathrm{I}]^2\kappa_3\kappa_{10}\kappa_{12}\kappa_{14}\kappa_{15}\kappa_{17} \\
p_2(\kappa,[\mathrm{I}]) = {} & \kappa_6\kappa_9\kappa_{11}\kappa_{13}\kappa_{17}+2\kappa_6\kappa_9\kappa_{11}\kappa_{13}\kappa_{18}+\kappa_6\kappa_9\kappa_{11}\kappa_{15}\kappa_{17}+2\kappa_6\kappa_9\kappa_{11}\kappa_{15}\kappa_{18}+\kappa_6\kappa_9\kappa_{13}\kappa_{14}\kappa_{17}+2\kappa_6\kappa_9\kappa_{13}\kappa_{14}\kappa_{18}+\kappa_6\kappa_9\kappa_{14}\kappa_{15}\kappa_{17} \\
& +2\kappa_6\kappa_9\kappa_{14}\kappa_{15}\kappa_{18}+[\mathrm{I}]^2\kappa_{10}\kappa_{12}\kappa_{14}\kappa_{15}\kappa_{17}+[\mathrm{I}]\kappa_6\kappa_{11}\kappa_{12}\kappa_{15}\kappa_{17}+[\mathrm{I}]\kappa_6\kappa_{11}\kappa_{12}\kappa_{15}\kappa_{18}+[\mathrm{I}]\kappa_9\kappa_{10}\kappa_{13}\kappa_{14}\kappa_{17}+[\mathrm{I}]\kappa_6\kappa_{12}\kappa_{14}\kappa_{15}\kappa_{17} \\
& +[\mathrm{I}]\kappa_9\kappa_{10}\kappa_{13}\kappa_{14}\kappa_{18}+[\mathrm{I}]\kappa_6\kappa_{12}\kappa_{14}\kappa_{15}\kappa_{18}+[\mathrm{I}]\kappa_9\kappa_{10}\kappa_{14}\kappa_{15}\kappa_{17} \\
p_3(\kappa,[\mathrm{I}]) = {} & \kappa_6\kappa_9\kappa_{11}\kappa_{13}\kappa_{18}+\kappa_6\kappa_9\kappa_{11}\kappa_{15}\kappa_{18}+\kappa_6\kappa_9\kappa_{13}\kappa_{14}\kappa_{18}+\kappa_6\kappa_9\kappa_{14}\kappa_{15}\kappa_{18}+[\mathrm{I}]^2\kappa_{10}\kappa_{12}\kappa_{14}\kappa_{15}\kappa_{17}+[\mathrm{I}]\kappa_6\kappa_{11}\kappa_{12}\kappa_{15}\kappa_{17}+[\mathrm{I}]\kappa_6\kappa_{11}\kappa_{12}\kappa_{15}\kappa_{18} \\
& +[\mathrm{I}]\kappa_6\kappa_{12}\kappa_{14}\kappa_{15}\kappa_{17}+[\mathrm{I}]\kappa_6\kappa_{12}\kappa_{14}\kappa_{15}\kappa_{18}-[\mathrm{I}]\kappa_9\kappa_{10}\kappa_{14}\kappa_{15}\kappa_{18} \\
p_4(\kappa,[\mathrm{I}]) = {} & \kappa_6\kappa_9\kappa_{11}\kappa_{13}\kappa_{18}+\kappa_6\kappa_9\kappa_{11}\kappa_{15}\kappa_{17}+2\kappa_6\kappa_9\kappa_{11}\kappa_{15}\kappa_{18}+\kappa_6\kappa_9\kappa_{13}\kappa_{14}\kappa_{18}+\kappa_6\kappa_9\kappa_{14}\kappa_{15}\kappa_{17}+2\kappa_6\kappa_9\kappa_{14}\kappa_{15}\kappa_{18}+[\mathrm{I}]^2\kappa_{10}\kappa_{12}\kappa_{14}\kappa_{15}\kappa_{17} \\
& +[\mathrm{I}]\kappa_6\kappa_{11}\kappa_{12}\kappa_{15}\kappa_{17}+[\mathrm{I}]\kappa_6\kappa_{11}\kappa_{12}\kappa_{15}\kappa_{18}+[\mathrm{I}]\kappa_6\kappa_{12}\kappa_{14}\kappa_{15}\kappa_{17}+[\mathrm{I}]\kappa_6\kappa_{12}\kappa_{14}\kappa_{15}\kappa_{18}+[\mathrm{I}]\kappa_9\kappa_{10}\kappa_{14}\kappa_{15}\kappa_{17}
\end{aligned}
$$

Figure 9: $\ker \Psi_{19}(\kappa)$ for the mass-action system in Figure 6.
